# Liver cancer initiation is dependent on metabolic zonation but decoupled from premalignant clonal expansion

**DOI:** 10.1101/2024.01.10.575013

**Authors:** Andrew Chung, Jason Guo, Yunguan Wang, Yuemeng Jia, Natasha Corbitt, Lin Li, Yonglong Wei, Min Zhu, Zixi Wang, Holly Guo, Purva Gopal, Guanghua Xiao, Tao Wang, Hao Zhu

**Author notes:** **Equal contributors** (Andrew Chung, Jason Guo). **Correspondence and lead contact:** Hao Zhu.

## Abstract

The origin of cancer is poorly understood because cells that obtain truncal mutations are rarely fate mapped in their native environments. A defining feature of the liver is zonation, or the compartmentalization of metabolic functions in hepatocytes located in distinct regions of the lobule ^1^. However, it is unknown if cancers develop in some zones but not others, and if there are metabolic determinants of cancer risk that track with cellular position. To study cancer initiation, we examined the effect of activating mutations in *Ctnnb1* and loss of function mutations in *Arid2*, two of the most commonly co-mutated genes in hepatocellular carcinoma (HCC) ^2^. We exploited glutamine synthetase (GS) as a faithful fate mapping marker of *Ctnnb1* mutant hepatocytes. By introducing mutations in distinct zones in a mosaic fashion, we showed that position and metabolic context regulate clone expansion. Mutant clones were maintained in zone 1 but largely outcompeted in zone 3. Paradoxically, clonal maintenance was anti-correlated with cancer initiation, as zone 3 mutant livers showed increased tumorigenesis. To define mechanisms, we individually deleted eleven zone specific genes in HCC mouse models, revealing that *Gstm2* and *Gstm3* were required for efficient HCC initiation in zone 3. These data indicate that liver cancer initiation is dependent on zonation but independent of clonal expansion.

---

Studying pre-malignant cells in their physiological environments would allow us to understand the competitive interactions between wild-type (WT) and mutant cells ^3^, the spatial metabolic factors that regulate transformation ^4,5^, and the cancer vulnerabilities associated with specific cells of origin ^6^. In the liver, chronic injury from viruses, alcohol, or overnutrition leads to cell death, regeneration, inflammation, fibrosis, cirrhosis, and ultimately, HCC ^7,8^. After decades of injury, cirrhotic livers consist of millions of regenerative nodules, most of which harbor somatic mutations ^9–11^. The relative infrequency of HCC development in this context raises questions about what happens to non-malignant clones that carry cancer mutations, and what factors regulate their fates. Sequencing of non-malignant tissues has exposed our incomplete understanding of premalignancy. Studies in the skin, lung, endometrium, esophagus, and cirrhotic liver have uncovered a broad array of somatic mutations ^9,10,12–15^. These studies suggest that the classical view of stepwise clonal expansions associated with newly acquired cancer mutations is not well understood, especially in solid organs. For example, *NOTCH* mutations are far more common in normal esophagus than in esophageal squamous cell carcinomas, suggesting that *NOTCH* mutant clones can expand but are protected from tumorigenesis ^15,16^. In contrast, several of the most frequently mutated genes in HCC, such as *TP53* and *CTNNB1*, are not frequently observed in the cirrhotic liver ^9–11^, suggesting that these mutations do not drive extensive clonal expansion events prior to tumor initiation. Altogether, the degree to which non-transformed cells harboring oncogenic driver mutations ultimately develop into cancer, or even clonally expand, is unknown.

In addition to a variety of genetic alterations, there are numerous liver cell types in which mutations can arise. Even among hepatocytes, there is extensive functional diversity associated with unclear malignant potential. Along the central-to-portal axis within the lobule, there are profound differences in metabolic gene expression, metabolic specialization, and oxygenation, collectively defined as “metabolic zonation” ^1,17^. Hepatocytes from different zones differ in their proliferative capacity ^18–20^, perhaps contributing to heterogeneous neoplastic potential. Because a majority of murine liver cancer models are not genetically or physiologically faithful to human disease, these models cannot be used to identify the cell type of origin or to track the evolution of HCCs. While previous studies have found that cells in zone 3 can transform into HCC ^21–23^, there has never been a head to head comparison to assess cancer formation from different zones. The recent generation of liver *CreER* mice with recombinase activity in distinct zonal populations permitted us to induce the same truncal HCC mutations in a variety of normal cell types ^19^. Using these tools, we 1) developed physiologically-relevant HCC mouse models, 2) elucidated how cellular position regulates mutant clone expansion and cancer risk, and 3) revealed genetic vulnerabilities associated with the metabolic zone of origin.

## Visual tracking of Ctnnb1 mutant cells in distinct hepatic zones

We focused on *CTNNB1* because it is one of the three most frequently mutated genes in HCC (mutation frequency based on TCGA: *TERT* promoter (44%), *TP53* (31%), *CTNNB1* (27%)). Given that *CTNNB1* mutations on their own are considered to be insufficient to drive cancer ^24^, we sought to co-mutate a second gene that would promote cancer development. We queried 14 genes that are either frequently mutated (*TERT, TP53, CTNNB1, ALB, APOB, AXIN1, ARID1A, ARID2*) or copy number altered in HCC (*NCOR1, RB1, CDKN2A, ERRFI1, PTEN, CCND1*)^2^. *CTNNB1* mutations were most likely to co-occur with *ARID2* loss-of-function mutations (**Fig. 1a**). ARID2, a tumor suppressor, is a component of the PBAF complex within the SWI/SNF family of ATP-dependent chromatin remodeling genes ^25^. Thus, we activated *Ctnnb1* and deleted *Arid2* in mouse livers. To maintain physiological gene regulation without frank overexpression, we used Cre-mediated recombination of endogenous *Ctnnb1* and *Arid2* floxed loci. We used the *ApoC4-CreERT2* mouse line to test mosaic recombination in hepatocytes across the entire liver lobule. Importantly, tamoxifen induction of CreER allowed for transient recombination of flox alleles at a specified time point, and avoided the persistent induction of mutations that would make fate mapping impossible. *ApoC4-CreERT2; Ctnnb1^(ex3)fl/+^; Arid2^fl/fl^; Rosa26^LSL-tdTomato/+^* mice were given tamoxifen at 4 weeks of age and sacrificed at 8 weeks (**Fig. 1b**). An ideal fate mapping marker of an oncogenically activated cell would be tightly linked to the oncogene being induced. In the liver, *Ctnnb1* activation faithfully induces ectopic glutamine synthetase (GS) expression outside of the endogenous expression normally observed in a 2-3 cell ring of hepatocytes around the central vein (CV) (**Fig. 1c, d**). Notably, GS expression is usually restricted to the peri-central region and not normally turned on during injury or regeneration. As expected, control mice (*Ctnnb1^(ex3)^ ^fl/+^; Arid2^fl/fl^* or *ApoC4-CreERT2; Arid2^fl/fl^*mice) only showed GS expression in peri-central hepatocytes (**Fig. 1c**). Thus, we used GS to lineage trace *Ctnnb1* mutant hepatocytes over space and time.

**Fig. 1.**
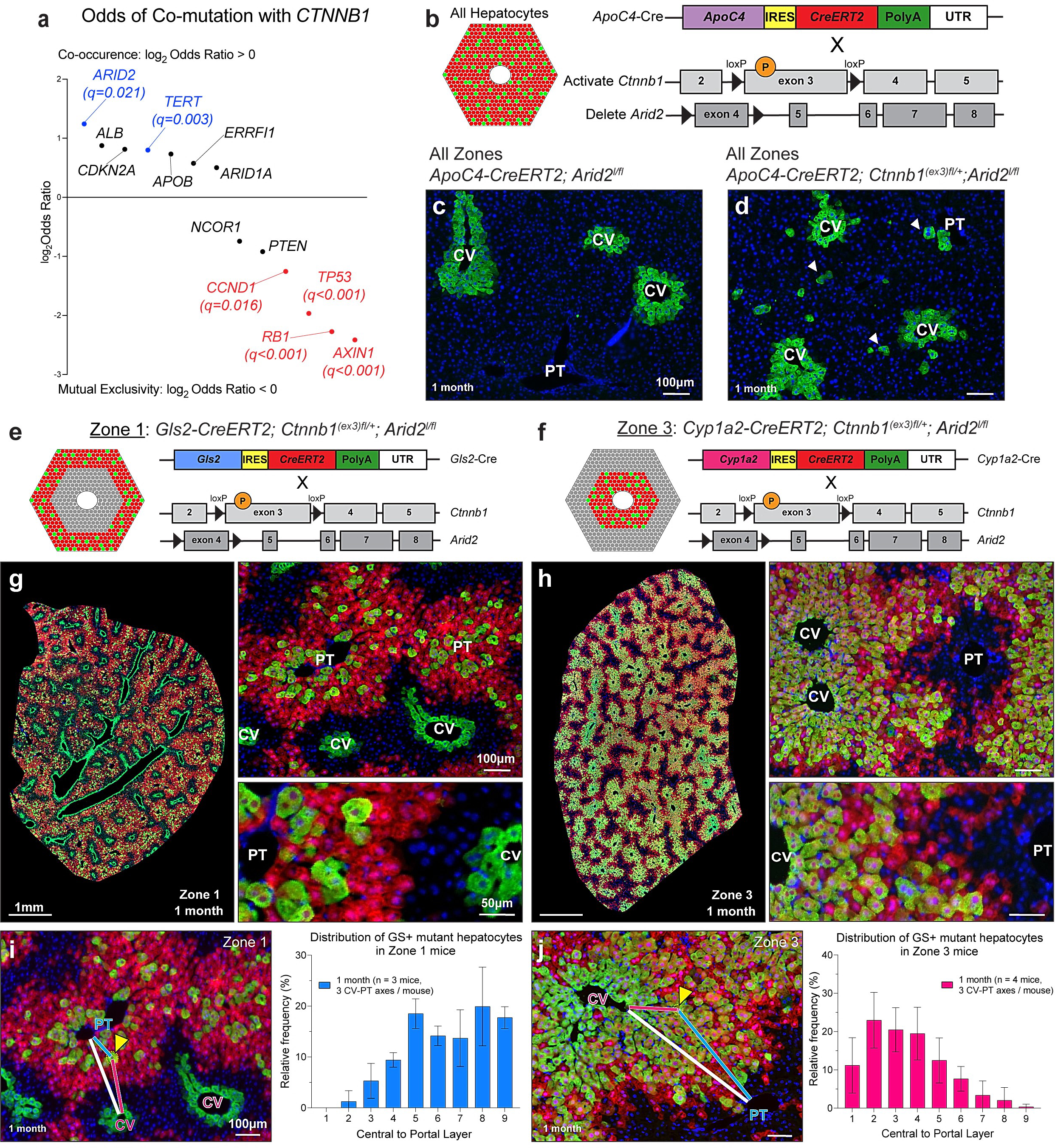
*Ctnnb1* mutant hepatocytes can be fate mapped with ectopic GS expression. **a.** Analysis of human HCC exome sequencing data from TCGA using log_2_ Odds Ratio. Genes with co-mutation q-values < 0.05 are highlighted for co-occurence (blue) or mutual exclusivity (red). **b.** Experimental scheme: *ApoC4-CreERT2; Ctnnb1^(ex3)^ ^fl/+^; Arid2^fl/fl^* mice that express *CreER* in all hepatocytes were injected with 100 mg/kg tamoxifen for 2 days (p28, p29) and analyzed 1 month later. **c.** *ApoC4-CreERT2; Arid2^fl/fl^* liver shows GS expression restricted around central veins (CV). **d.** *ApoC4-CreERT2; Ctnnb1^(ex3)/fl/+^; Arid2^fl/fl^* liver shows ectopic GS-expressing mutant hepatocytes (arrowheads) scattered throughout the lobule. **e, f.** Experimental scheme: Zone 1 *Gls2-CreERT2; Ctnnb1^(ex3)^ ^fl/+^; Arid2^fl/fl^* mice (**e**) or Zone 3 *Cyp1a2-CreERT2; Ctnnb1^(ex3)^ ^fl/+^; Arid2^fl/fl^* mice (**f**) were injected with 100 mg/kg tamoxifen for two days (p28, p29) and sacrificed at later time points for analysis. **g.** Zone 1 *Gls2-CreERT2; Ctnnb1^(ex3)^ ^fl/+^; Arid2^fl/fl^; Rosa26^LSL-tdTomato/+^* liver immunostained for GS at 1 month post-tamoxifen shows localization of mutant hepatocytes. **h.** Zone 3 *Cyp1a2-CreERT2; Ctnnb1^(ex3)^ ^fl/+^; Arid2^fl/fl^; Rosa26^LSL-tdTomato/+^* liver immunostained for GS 1 month post-tamoxifen shows localization of mutant hepatocytes. **i.** Positional index (P.I.) was calculated by measuring the distances from a GS+ cell (yellow arrowhead) to the nearest CV (red line), the nearest PT (blue line), and the distance between the CV and PT (white line). Shown are PI measurements in a zone 1 liver 1 month post-tamoxifen. The distribution of GS+ mutants is calculated by relative frequency of GS+ cells in zone 1 (layers 7-9), zone 2 (layers 4-6), and zone 3 (layers 1-3). **j.** P.I. distance measurements for a single GS+ cell in a representative zone 3 mutant liver. On the right, distribution of GS+ mutants by layers at 1 month post-tamoxifen. In all P.I. measurements, the n for individual mice is denoted in the graph. At least three PV-CV axes are quantified for each mouse. Data in bar graphs are displayed as mean ± SEM.

Given the large differences in zonal gene expression across the lobule, we reasoned that the hepatocyte subtype could have a major effect on clone fates. To trace mutant clones in different zones, we generated *Gls2-CreERT2; Ctnnb1^(ex3)fl/+^; Arid2^fl/fl^; Rosa26^LSL-tdTomato/+^*mice (zone 1 mutant mice, **Fig. 1e**) and *Cyp1a2-CreERT2; Ctnnb1^(ex3)^ ^fl/+^; Arid2^fl/fl^; Rosa26^LSL-tdTomato/+^* mice (zone 3 mutant mice, **Fig. 1f**). Tomato reporter expression confirmed that CreER recombination was restricted to hepatocytes around portal triads (PT) in zone 1 mutant mice (**Fig. 1g**) and around CVs in zone 3 mutant mice (**Fig. 1h**). In both types of mutant mice, only a subset of Tomato+ hepatocytes were also GS+, indicating that CreER-mediated recombination occurred in a mosaic fashion. While this indicated that CreER was not 100% efficient, it also faithfully modeled the somatic mosaicism seen in human tissues prior to cancer development. To gain a quantitative view of spatial distribution of mutant hepatocytes within the lobule, we measured the zonal position index (P.I.) of mutant hepatocytes in livers from zonal mice (method described in ^26^). The P.I. provided a continuous readout of zonal position that can be binned into 9 layers, based on a previously established convention with layer 1 being the most central and 9 being the most portal ^27^. In the zone 1 mice, mutant hepatocytes were initially in zones 1 and 2, with highest frequency in layers 5-9 (**Fig. 1i**). In the zone 3 mice, mutant hepatocytes were initially in zones 2 and 3, with highest frequency in layers 2-5 (**Fig. 1j**). This confirmed that mutations were induced in a zone-specific manner.

## Ctnnb1 mutant cells outcompete WT cells in zone 1 but not in zone 3

To determine if *Ctnnb1/Arid2* mutant hepatocytes originating in zone 1 and 3 exhibit differences in clonal fates, we lineage-traced mutant hepatocytes. After induction of mutations at 1 month of age, livers were analyzed 1, 2, 3, and 6 months later. The frequency of GS+ mutant hepatocytes in zone 1 mutant mice was stable over time (**Fig. 2a**). In contrast, the frequency of mutant hepatocytes in zone 3 mutant mice decreased over time (**Fig. 2b**). 6 months after tamoxifen, GS+ cells in zone 1 mutant mice clustered around portal tracts, but in zone 3 mutant mice, ectopic GS+ cells disappeared such that most of the GS+ cells remaining were pericentral hepatocytes with endogenous GS expression (**Fig. 2c**). To determine if the remaining GS+ cells at the 6 month time point were mutant or WT, we used IHC to determine if CTNNB1 was membrane bound or cytoplasmic/nuclear, which would be indicative of activated mutant CTNNB1. The cells with nuclear CTNNB1 mirrored the frequency and localization of ectopic GS in both zone 1 and zone 3 mutant mice (**Fig. 2d**).

**Fig. 2.**
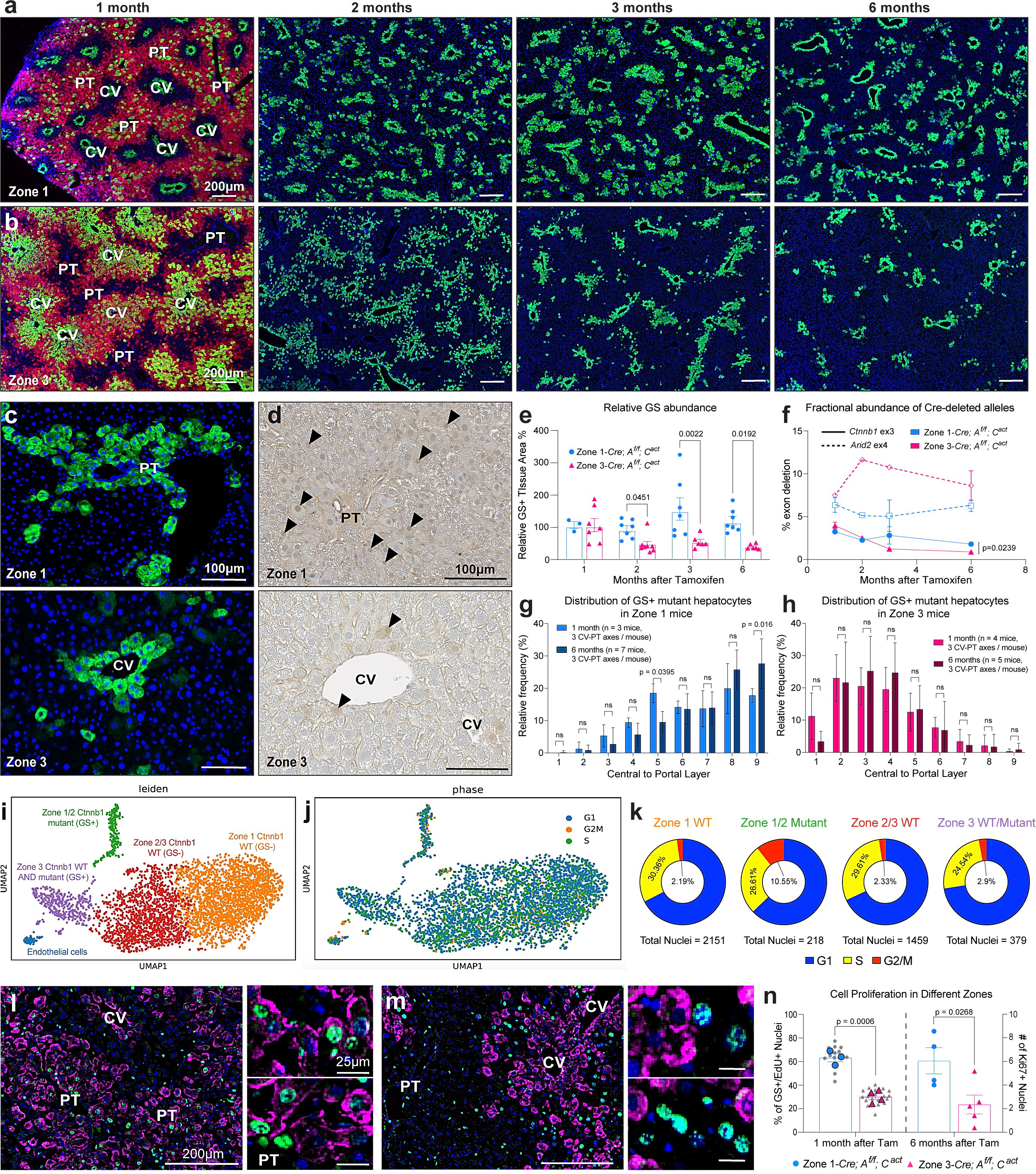
*Ctnnb1* mutant clones are maintained in zone 1 but disappear from zone 3. **a.** GS-stained livers from zone 1 *Gls2-CreERT2; Ctnnb1^(ex3)^ ^fl/+^; Arid2^fl/fl^* mice from 1, 2, 3, and 6 months after tamoxifen. **b.** GS-stained livers from zone 3 *Cyp1a2-CreERT2; Ctnnb1^(ex3)^ ^fl/+^; Arid2^fl/fl^* mice from 1, 2, 3, and 6 months after tamoxifen. **c.** Portal triad (PT) region of GS-stained liver from zone 1 mutant mice (top panel) and central vein (CV) region of liver from zone 3 mutant mice (bottom panel) at 6 months post-tamoxifen. Zone 1 mutants persist over time and zone 3 mutants are reduced in number. **d.** IHC staining for CTNNB1 in zone 1 (top) and zone 3 (bottom) mutant mice at 6 months after tamoxifen shows the persistence of mutant clones (arrowheads) that have increased nuclear and cytoplasmic CTNNB1 staining in zone 1 compared to the loss of mutants and more membranous staining in zone 3. **e.** Image analysis was used to calculate the GS+ area in zone 1 (blue) and zone 3 (red) mice at 1, 2, 3, and 6 months post-tamoxifen. GS+ area percentages for each zone mouse type were normalized to the 1 month time point, which was designated as 100%. For zone 1 mice at 1, 2, 3, and 6 months, each dot represents one mouse and n = 3, 7, 7, 7 mice. For zone 3 mice at 1, 2, 3, and 6 months, each dot represents one mouse and n = 7, 7, 6, 6 mice. **f.** ddPCR was used to measure the abundance of *Cre*-deleted alleles for *Ctnnb1* exon 3 (solid lines) and *Arid2* exon 4 (dashed lines) in zone 1 (blue) and zone 3 (red) mice at 1, 2, 3, and 6 months post-tamoxifen. Allele percentages for each gene in each zonal mouse type are shown. For zone 1 mice at 1, 2, 3, and 6 months: n = 2, 1, 2, 7, respectively. For zone 3 mice at 1, 2, 3, and 6 months: n = 2, 1, 1, 5, respectively. **g.** Distribution of GS+ hepatocytes in lobular layers at 1 and 6 months post-tamoxifen as measured by P.I. This shows a relative shift toward the PT within zone 1 livers, with a decrease in midlobular layer 5 and increase in periportal layer 9. **h.** Distribution of GS+ hepatocytes in lobular layers at 1 and 6 months post-tamoxifen as measured by P.I. This shows no shift within zone 3 livers. In all P.I. measurements, the n for individual mice is denoted in the graph. At least three PV-CV axes are quantified for each mouse. **i.** snRNA-seq of a zone 3 mutant liver identifies five cell populations. **j.** The cell cycle phase of every nucleus sequenced was predicted using reference cell cycle genes. **k.** The number of nuclei in G1, S, and G2/M (blue, yellow, and red) phases was estimated in the four major hepatocyte populations. More zone 1 and zone 2 mutant hepatocytes were in the G2/M phase relative to the other populations (Chi-square test, ****p < 0.0001). **l.** Zone 1 (n = 3) and zone 3 (n = 4) mice were given 10 days of EdU water starting 1 month post-tamoxifen. Zone 1 liver showing EdU+ and GS+; EdU+ hepatocytes 1 month after tamoxifen (left panel) with zoomed-in images showing nuclei around portal triad (PT, right panel). **m.** Zone 3 liver immunostained for GS and stained for EdU showing EdU+ and GS+; EdU+ hepatocytes 1 month after tamoxifen (left panel) with zoomed-in images showing nuclei around CV (right panel). **n.** The percentage of GS+/EdU+ cells over EdU+ cells were measured. Smaller black dots represent percentages from single 10X images, and larger colored dots represent percentages from each mouse averaged from 4 or more 10X images (left axis). EdU pulse chase experiments were performed two independent times and the data was aggregated. Additionally, zone 1 (n = 4 mice) and zone 3 (n = 5 mice) mutant livers were stained and quantified for Ki67 at 6 months post-tamoxifen (right axis). Data in bar graphs are displayed as mean ± SEM, and statistical analyses were performed using two-way ANOVA when comparing the two zone groups at multiple time points ( **e,f,g,h**) or a two-tailed, unpaired Student’s t-test when comparing two groups at a single time point (**n**).

To better quantify the abundance of mutant clones, we also implemented an image analysis algorithm (**Extended Data Fig. 1**). The relative image area occupied by GS+ cells was unchanged in the zone 1 mutant livers from 1 to 6 months, while the GS+ area in zone 3 mutant livers decreased (**Fig. 2e**). The relative GS+ area normalized to the 1 month time point was 3-fold greater in zone 1 vs. 3 mutant livers at 6 months. To address the possibility that GS was not tracking all *Ctnnb1* mutant clones, we also quantified mutations using an orthogonal digital droplet PCR (ddPCR) assay (**Extended Data Fig. 2**). In zone 1 mutant mice, the relative fraction of mutant *Ctnnb1* alleles at 6 months normalized to the 1 month time point was 55.9%, while in the zone 3 mutant mice, the relative fraction was 22.5% (p = 0.0589, **Fig. 2f**). Altogether, these results showed that *Ctnnb1* mutant hepatocytes in zone 3 decline in number to a greater degree than those in zone 1.

Having observed substantial differences in mutant clone fates in zone 1 and 3, we measured changes in the zonal position of mutant hepatocytes over time (**Extended Data Fig. 3**). In zone 1 mutant mice, there was a shift in position of mutant hepatocytes towards the portal tracts such that the proportion of hepatocytes in midlobular layer 5 decreased, while the proportion of hepatocytes in the most portal layer 9 increased after 6 months (**Fig. 2g**). In zone 3 mutant mice, the decline in GS+ cells was not characterized by a positional shift, indicating that mutant clones were lost throughout zone 3 (**Fig. 2h**). The relative shift of mutant hepatocytes in the zone 1 mice is occurring in the context of mutant clone stability, suggesting that mutant hepatocytes located in the most periportal layers proliferated to maintain the overall mutant clone abundance. This further supports the observation that zone of origin is a critical determinant of mutant hepatocyte fate.

## Characterization of mutant livers with single nuclear sequencing

To further explore hepatocytes in the zone 3 mutant liver, we used single nuclear RNA sequencing (snRNA-seq). SnRNA-seq identified four unique hepatocyte populations (**Fig. 2i**). Zone 1 hepatocytes expressed periportal genes such as *Sds, Hal, Gls2,* and *Ass1* (orange dots in **Fig. 2i**; **Extended Data Fig. 4** and **Extended Data Table 1**) ^17,27^. Zone 2 and 3 hepatocytes were identifiable by their enriched expression of genes known to peak in zone 2 (*Hamp2, Gpam, Cyp3a25*) and 3 (*Akr1c6, Plpp3, Cyp27a1*) (red dots in **Fig. 2i**; **Extended Data Fig. 4** and **Extended Data Table 2**). These two populations were predominantly *Ctnnb1* WT because they did not express *Glul* (GS). Zone 3 hepatocytes expressed pericentral genes such as *Glul (GS), Slc1a2, Cyp1a2, Cyp2e1,* and *Lgr5* (purple dots in **Fig. 2i**; **Extended Data Fig. 4** and **Extended Data Table 3**). Given that these nuclei were from a zone 3 mutant mouse, this population is likely a mixture of *Ctnnb1* WT and mutant hepatocytes. Finally, a population of hepatocytes simultaneously expressing zone 1 and 3 genes (*Gls2, Glul (GS), Cyp1a2*) were likely to be *Ctnnb1* mutant cells positioned within zones 1 or 2 (green dots in **Fig. 2i**; **Extended Data Fig. 4** and **Extended Data Table 4**). In summary, we identified zone 1 WT, zone 2/3 WT, zone 3 WT and mutant, and zone 1/2 mutant hepatocytes. To characterize the behavior of these populations, we examined cell cycle phase-specific gene expression (**Fig. 2j**). A higher proportion of the zone 1/2 mutant hepatocytes were in the G2/M cell cycle phase (**Fig. 2k**), suggesting that these hepatocytes were more proliferative than other hepatocyte populations.

## The relative fitness of mutant hepatocytes depends on their zonal location

To functionally explore why mutants were more competitive in zone 1 than they were in zone 3, we assessed liver cell proliferation. After a 10 day EdU pulse chase, we observed an increased absolute number of proliferating hepatocytes in zone 1 mutant vs. WT livers at the same age (**Extended Data Fig. 5a,b**). Among the EdU+ hepatocytes in zone 1 mutant mice, a majority (65%) were GS+ Ctnnb1 mutants. Thus, EdU+ cells were more frequently located in zone 1 because the GS+ clones were predominantly in zone 1 (**Fig. 2l,n**). In zone 3 mutant mice, there was a trend toward an increased absolute number of proliferating hepatocytes as compared to WT mice (**Extended Data Fig. 5a,b**). However, among the EdU+ hepatocytes in zone 3 mutant mice, only 30% of EdU+ cells were GS+, thus most proliferating cells were WT (**Fig. 2m,n**). In agreement with EdU tracing, we also observed more Ki-67+ nuclei in the zone 1 mutant livers at the 6 month time point (**Fig. 2n, Extended Data Fig. 5c-n**).

To directly assess competition between WT and mutant cells within zones 3 mutant mice, we traced Tomato+; GS-WT cells and Tomato+; GS+ mutant cells. After 6 months of lineage tracing, the tdTomato+; GS-WT population expanded relative to the tdTomato+; GS+ mutant population, indicating that WT cells from within zones 2 and 3 were outcompeting mutant cells (**Extended Data Fig. 6a**). These data suggest that in zone 1, mutant GS+ hepatocytes were more competitive than WT hepatocytes due to increased proliferation. In zone 3, mutant GS+ hepatocytes were less competitive than WT hepatocytes, at least in part due to reduced proliferation. Taken together, the relative fitness of mutant hepatocytes depends on both zone of origin and mutation status.

## Ctnnb1 and Arid2 mutations drive HCC development in a synergistic fashion

Our next objective was to quantify the malignant fates of these pre-malignant clones. A fundamental problem in liver cancer modeling is that most mouse models of HCC development require non-physiologic levels of oncogene overexpression or chemical mutagenesis. Models involving *Myc* overexpression, *Pten* deletion, or DEN mutagenesis are less clinically relevant and do not involve the most common HCC driver mutations. We first asked if *Ctnnb1* or *Arid2* mutations are sufficient to drive liver cancers in mouse models. Few mice from any of the *Ctnnb1* or *Arid2* models developed macroscopic tumors prior to the six month time point, thus we performed cancer analysis in mice that were aged for 6-12 months (**Fig. 3a,b**). In accord with the modest rate of cancer development in patients with cirrhosis, our data suggests that these two mutations drive cancer with a similar kind of slow kinetics as seen in human liver disease. This is in contrast to overexpression or hydrodynamic transfection models that cause rapid cancer development within 1-3 months. Even after 6 months, mice without CreER did not develop any observable tumors (**Fig. 3c**). In either zone, only 1 of 41 *Arid2* single mutant mice and 4/12 *Ctnnb1* single mutant mice developed unifocal lesions, indicating that *Arid2* mutations alone or *Ctnnb1* mutations alone were inefficient for carcinogenesis (**Fig. 3c**). In contrast, 40/50 (80%) double mutant mice with mutations in either zone developed cancers, usually in a multifocal pattern (**Fig. 3a-c**). As suggested by the TCGA analysis, co-mutating *Ctnnb1* and *Arid2* resulted in efficient liver tumor formation.

**Fig. 3.**
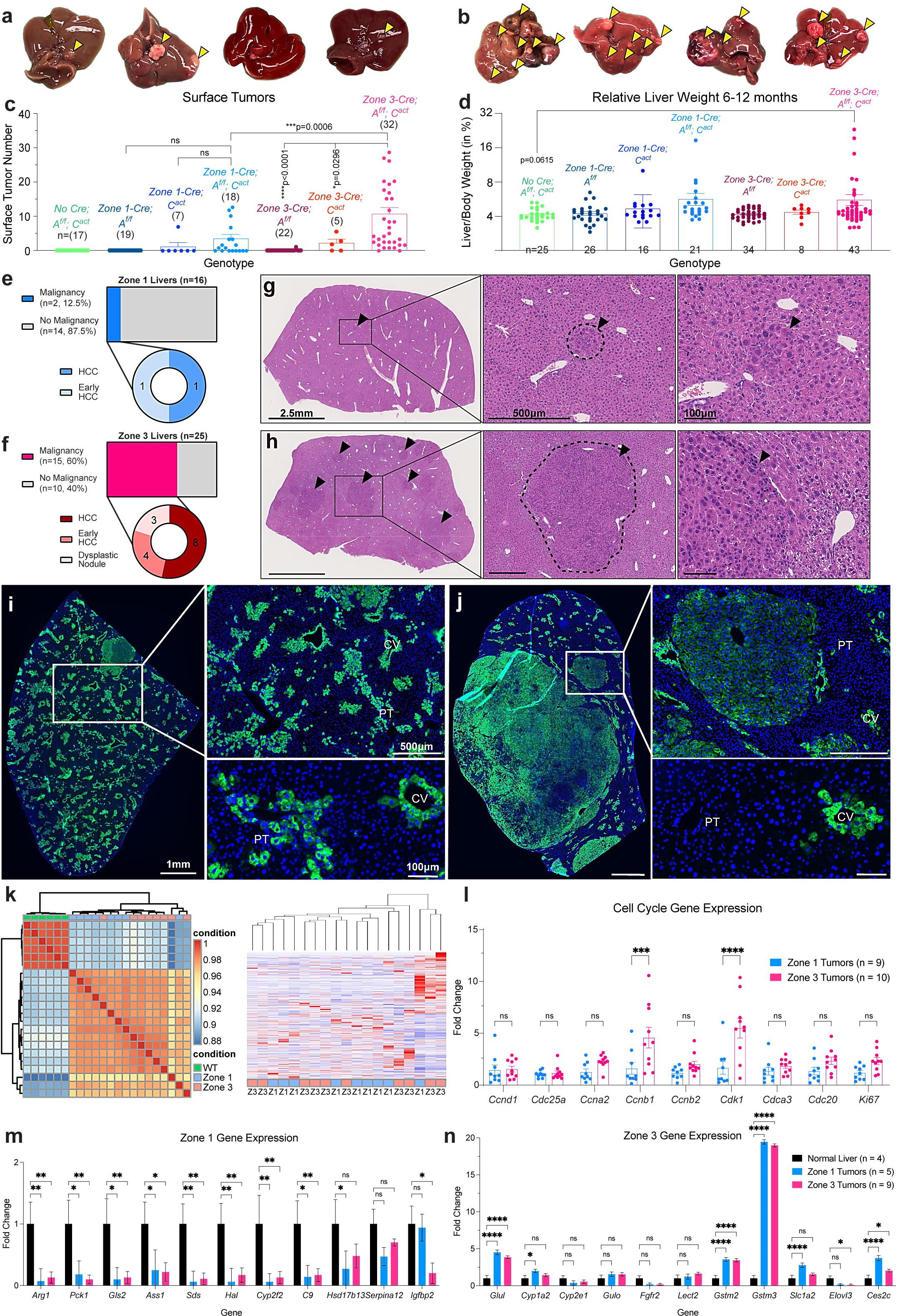
Pre-malignant clonal expansion is anti-correlated with the frequency of cancer initiation. **a,b.** Zone 1 *Gls2-CreERT2; Ctnnb1^(ex3)^ ^fl/+^; Arid2^fl/fl^* and zone 3 *Cyp1a2-CreERT2; Ctnnb1^(ex3)^ ^fl/+^; Arid2^fl/fl^* livers were collected 6-12 months after tamoxifen, and gross images of livers are shown. Arrowheads point to surface tumors. **c.** Surface tumors in livers collected 6-12 months post-tamoxifen were quantified. Zone 3 mice had significantly more liver tumors than zone 1 mice (median 5.5 vs. 1, p = 0.02126; mean 10.72 vs. 3.50, p = 0.0006). Each dot is an individual mouse, n is indicated in parentheses. **d.** Liver to body weight ratios (%) were used as a surrogate metric for tumor burden in livers collected 6-12 months post-tamoxifen. Each dot is an individual mouse, n is indicated at the base of each bar. **e,f.** Blinded pathological analysis of histology for zone 1 (**e**) and zone 3 (**f**) mice identified the presence of nodules and classified these nodules as HCC, early HCC, or dysplastic nodules, and the resulting percentages are shown (Chi-square test, *p=0.035). **g,h.** Whole liver lobe histology of a zone 1 liver at 11 months **(g)** and a zone 3 liver at 10 months **(h)**. Arrowheads point to HCCs. Zoomed-in images (center, right panels) focus on tumor histology. Zone 1 tumors are less frequent and smaller than zone 3 tumors. **i.** Zone 1 liver immunostained for GS 8 months post-tamoxifen shows fewer tumors. Zoomed-in images (right panels) highlight a non-malignant GS+ mutant population that is maintained around the portal tracts along the portal-to-central vein axis. **j.** Zone 3 liver immunostained for GS 8 months post-tamoxifen shows multiple tumors of various sizes. Zoomed-in images (right panels) highlight a loss of GS+ mutant cells around the CV. **k.** Unsupervised clustering of bulk RNA-seq in tumors from zone 1 (n = 9) and zone 3 (n = 10) cluster together and are not distinct, indicating that these tumors are transcriptionally similar. **l.** RT-qPCR for cell cycle genes reveals that zone 3 tumors upregulate certain cell cycle genes, particularly those related to G2-M transition, compared to zone 1 tumors **m.** RT-qPCR for zone 1 genes reveals that tumors originating in both zones downregulate most zone 1 genes compared to normal liver. **n.** RT-qPCR for zone 3 genes reveals that tumors originating in both zones upregulate a subset of zone 3 genes compared to normal liver. Data in bar graphs are displayed as mean ± SEM, and statistical analyses were performed using a one-way ANOVA ( **c,d**), Fisher’s exact test (**e,f**), or two-way ANOVA ( **l,m,n**). Significance is displayed as ****p < 0.0001, ***p < 0.001, **p < 0.01, *p < 0.05, ns = not significant.

## Pre-malignant clonal dynamics anti-correlate with tumorigenic potential

Since mutant hepatocyte clones were more competitive in zone 1, we hypothesized that zone 1 mice would develop more tumors than zone 3 mice. Surprisingly, 30/32 (94%) of zone 3 mice developed visible surface tumors while only 10/18 (56%) of zone 1 mice developed tumors (**Fig. 3c**). Also, zone 3 livers harbored a higher number of surface tumors (median: 5.5 vs. 1, p = 0.02126; mean: 10.72 vs. 3.50, p = 0.0006) (**Fig. 3c**). Liver to body weight ratios, a surrogate measure for tumor development, followed a similar pattern (**Fig. 3d**). Because not all liver lesions are malignant and not all malignancies are HCC, we performed blinded histologic analysis of microscopic tumors. This showed that 16/16 nodules from 25 zone 3 mice were malignant (dysplastic nodules, early HCCs, or HCCs), whereas only 2/6 nodules that arose from the 16 zone 1 mice were malignant, and the 4/6 remaining nodules had uncertain pathologic significance (**Fig. 3e-h**). Overall, 16/25 (64%) of zone 3 mice and 2/16 (12.5%) of zone 1 mice harbored malignant lesions. In 8 month-old mice that had formed tumors, GS staining once again showed a maintenance of mutant clones in zone 1 mice (**Fig. 3i**), whereas zone 3 mice showed almost no ectopic GS+ clones but a substantial increase in tumor burden (**Fig. 3i,j**). Furthermore, the tumors from either mice stained strongly for GS, confirming that these tumors are universally driven by oncogenic CTNNB1 (**Fig. 3i,j**).

Since differential efficiency of CreER activity in zone 1 and 3 mice could account for differences in HCC frequency, we performed experiments to interrogate this possibility. Three mutant populations needed to be considered for this question: *Arid2* single mutants, *Ctnnb1* single mutants, and *Ctnnb1; Arid2* double mutants. Because *Arid2* single mutant mice did not develop cancer at an appreciable rate (1/41 mice had a single gross lesion), we reasoned that this population of mutant hepatocytes did not require monitoring. *Ctnnb1* single mutant mice were able to generate HCC, so we performed lineage tracing of *Ctnnb1* single mutant cells to determine if clonal dynamics also reflected HCC risk in this setting. 2 of 5 mice (40%) with *Ctnnb1* single mutants in zone 3 developed tumors while only 1 of 7 mice (14%) with *Ctnnb1* single mutants in zone 1 developed tumors (**Fig. 3c**). The increased recombination efficiency in the *Cyp1a2-CreER* zone 3 mice could indeed cause a higher rate of HCC development in these mice. However, most GS+ clones also disappeared in zone 3 single mutant mice by the 6 month time point (**Extended Data Fig. 6b**). This indicated that even when all mutant hepatocytes potentially responsible for tumorigenesis can be visualized in a single mutant mouse, there is still discord between the number of mutant hepatocytes and the eventual number of tumors. Because the GS+ population also declines in a similar fashion in the *Ctnnb1; Arid2* zone 3 mice, it is likely that the double mutant population declines similarly. Since GS only allows tracking of *Ctnnb1* mutant cells without regard to *Arid2* mutation status, it is formally possible that the double mutant clones were maintained or expanded in zone 3. To address this, we approximated the number of double mutant cells using a ddPCR assay that probes for floxed deletions. At the 1 month time point, the double mutant rates were 0.21%±0.059 in zone 1 mutant mice and 0.29%±0.024 in zone 3 mutant mice (mean±SD; n = 2,2). At 6 months, the frequencies were 0.12%±0.059 in zone 1 mice vs. 0.078%±0.057 in zone 3 mice (mean±SD; n = 5,5). The frequency of double mutant cells at the 6 month time point trended lower in the zone 3 mice, an observation that is the opposite of the null hypothesis which expected more double mutants in zone 3. This indicates that differential Cre efficiency was not responsible for differences in tumorigenesis.

## Transcriptional characterization of tumors from distinct zones

We then asked if tumors from different zones showed gene expression differences that paralleled their respective zones of origin. We profiled both *Ctnnb1* single mutant and *Ctnnb1; Arid2* double mutant tumors from both zones (n = 9 from zone 1 and n = 10 from zone 3), and were surprised to find that the samples did not segregate according to zone of origin or genotype (**Fig. 3k** and **Extended Data Fig. 7a**). When we used GSEA to compare tumors from opposing zones, we found that tumors from zone 3 had higher cell cycle and lower metabolic gene expression (**Extended Data Fig. 7b** and **Fig. 3l**). To determine if tumors retained zone-specific metabolic gene expression, we examined periportal (zone 1) genes and pericentral (zone 3) genes. Compared to control liver tissues, all tumors downregulated periportal (zone 1) genes (**Fig. 3m**) and upregulated a subset of pericentral (zone 3) genes, including *Glul, Gstm2, Gstm3,* and *Ces2c* (**Fig. 3n**). While a small number of genes were differentially expressed between zone 1 and zone 3 tumors (**Extended Data Fig. 7c, Extended Data Table 5**), these tumors predominantly exhibited a zone 3 gene expression program consistent with the role of WNT activation in defining zone 3 identity.

## Integrated genomic and functional exploration of zonated genes that promote HCC

These results indicate that the clonal dynamics of mutant hepatocytes originating from different zones do not necessarily predict tumorigenic potential. Rather, other key zone-defining factors may be more influential for cancer initiation. In order to functionally identify genes that regulate the rate of mutant clone transformation, we integrated data from snRNA-seq of the liver and *in vivo* CRISPR screening ^9,28^. We focused our attention on ∼100 of the most zonated genes expressed in zone 1 and zone 3 ^17^. Because this was too many genes to functionally assess individually, we used *in vivo* CRISPR knockout (CRISPR KO) and CRISPR activation (CRISPRa) screens (**Extended Data Fig. 8a,b, Extended Data Table 6**) to narrow down the list to genes that influence hepatocyte proliferation in *Fah* KO mice. We identified genes that affected proliferation and survival by assessing changes in sgRNA abundance between the initial library and the regenerated livers after 4 weeks of hepatocyte repopulation (**Extended Data Fig. 8c,d**). We also used the bulk RNA-seq data from the *Ctnnb1; Arid2* HCC models to look for differences between tumors arising from zones 1 and 3. Altogether, we chose 11 genes from an integrative analysis of the CRISPR screens, the snRNA-seq, and the tumor RNA-seq as candidate factors that might regulate cancer development in different zones (**Fig. 4a**).

**Fig. 4.**
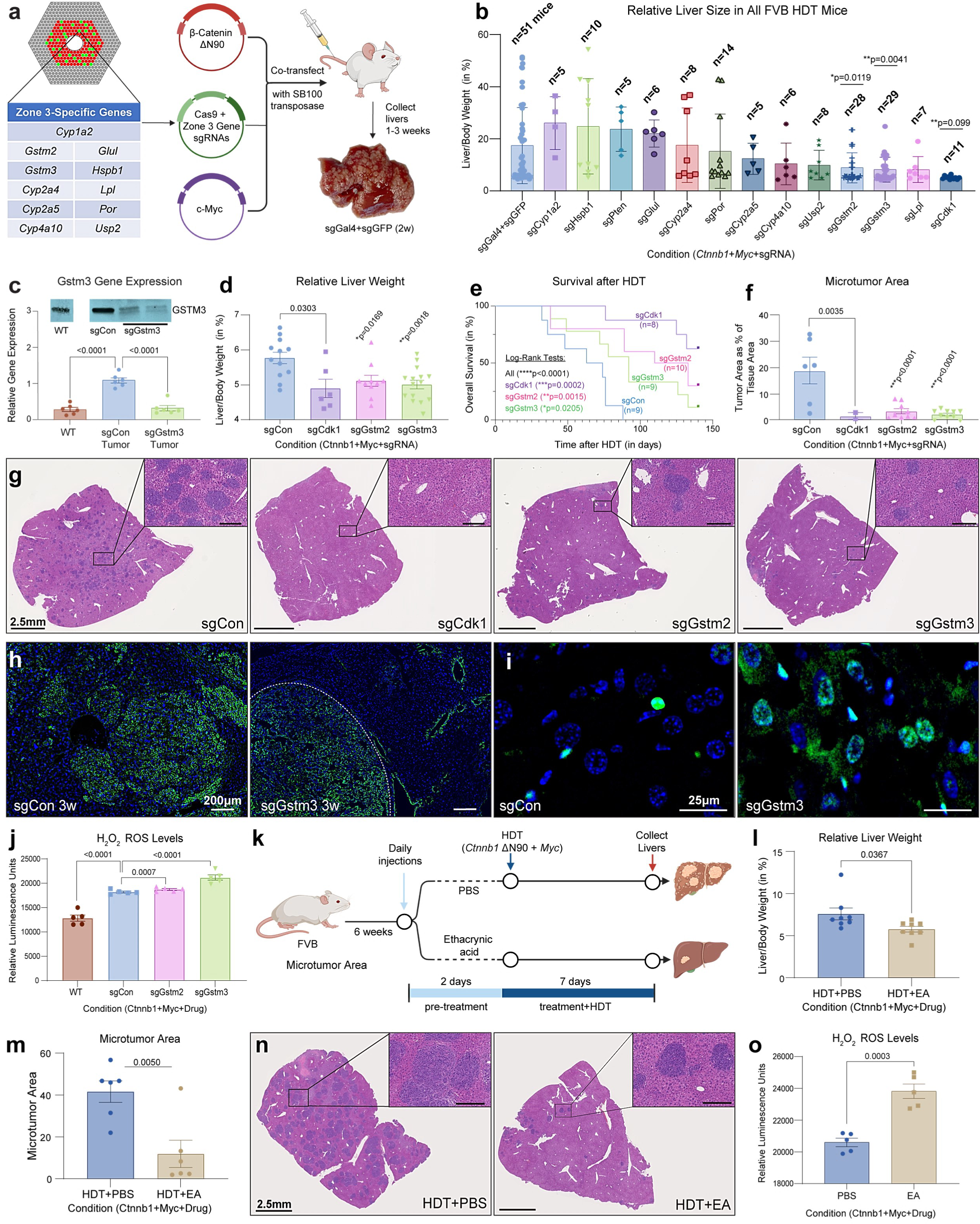
GSTM2 and GSTM3 are zonated metabolic enzymes that promote tumor initiation in zone 3. **a.** Experimental scheme: WT FVB mice were hydrodynamically transfected at 6 weeks of age with 2mL of the transposon and transposase plasmids and sacrificed 1-3 weeks post-injection. 11 genes of interest were chosen from an integrated analysis of snRNA-seq, bulk RNA-seq, and in vivo CRISPR screening data. **b.** Liver to body weight ratio (%) was used as a surrogate for tumor burden in HDT mice. Each dot is one mouse, and the number of mice used for each model is noted above the bars. **c.** RT-qPCR for *Gstm3* mRNA levels and western blot for GSTM3 protein levels. The mRNA fold expression is relative to sgGal4+sgGFP (together denoted as sgCon) livers, and each dot represents one liver sample from one mouse and n = 6 mice per group. Each western blot lane represents lysate from an individual mouse, and non-adjacent lanes from the same blot are separated. **d.** LW/BW at 1 week post-HDT in FVB mice for sgCon (n=12 mice), sgCdk1 (n=4), sgGstm2 (n=10), and sgGstm3 (n=15). Each dot represents one mouse. HDT experiments for these groups were performed three independent times, and data was aggregated. **e.** Kaplan-Maier curves for HDT injected mice. Survival was followed until death or for 150 days. The number of mice used in each group is noted in the figure. **f.** Quantification of microscopic tumor area relative to total tissue area for sgCon (n=6 mice), sgCdk1 (n=2), sgGstm2 (n=9), and sgGstm3 (n=10) livers collected 1 week after HDT. Each dot represents one liver from one mouse. **g.** H&E staining 1 week after HDT showing microscopic tumors that have high nuclear/cytoplasmic ratios. Insets are zoomed in for tumor nodules (inset scale bars = 250μm). **h.** sgCon (left panel) and sgGstm3 (right panel) liver sections were immunostained for GS at 3 weeks post-HDT, revealing more tumors in the control group (dotted circle). **i.** Apoptosis in sgCon (top panel) and sgGstm3 (bottom panel) livers assessed by TUNEL (green) staining 1 week post-HDT. **j.** ROS as measured by H_2_O_2_ luminescence levels in FVB WT (n=4 mice), sgCon (n=5), sgGstm2 (n=5), and sgGstm3 livers (n=5). Each dot represents one liver sample from one mouse. **k.** Experimental scheme: WT FVB mice were pretreated with PBS (vehicle) or 10 mg/kg ethacrynic acid in PBS (EA) for two days prior to HDT with *CTNNB1* + *MYC* and daily thereafter before sacrifice at 1 week post-HDT. **l.** LW/BW ratios at 1 week after HDT in PBS (n = 8) and EA (n = 8) groups. Each dot represents one mouse. Drug treatment HDT experiments were performed two independent times, and data was aggregated. **m.** Quantification of microtumor area for PBS (n = 6) and EA (n = 6) mice. Each dot represents one liver from one mouse. **n.** Representative H&E staining of control and EA-treated liver sections showing microtumors, with close up insets (inset scale bars = 250μm). **o.** ROS as measured by H_2_O_2_ luminescence levels in PBS (n=5) and EA (n=5) mice. Each dot represents one liver sample from one mouse. Data in bar graphs are displayed as mean ± SEM, and statistical analyses were performed using a one-way ANOVA when comparing multiple HDT groups ( **b,c,d,f,j**) or a two-tailed and unpaired Student’s t-test when comparing two groups (**l,m,o**). For survival analysis, a Log-rank Mantel-Cox test was performed to compare all groups, and individual tests were performed to compare sgCon to every other group, as indicated (**e**).

To explore the mechanistic basis by which zone 3 mutant hepatocytes have more tumorigenic potential, we deleted these 11 genes, with the expectation that their loss would influence tumor initiation. Using hydrodynamic transfection (HDT) to deliver Sleeping Beauty transposon plasmids, we generated mouse models with CRISPR-mediated deletion of a single gene of interest concurrent with overexpression of mutant *CTNNB1* and *MYC* (**Fig. 4a**). While delivery of either *CTNNB1* or *MYC* alone was insufficient to drive cancer within 4 weeks ^29^, delivering both led to liver carcinogenesis within 1-2 weeks on the FVB strain background. This rapid tumorigenesis, while less physiologic, provided a tractable system for identification of essential genes. sgRNA against *GAL4* or *GFP* were used as negative controls (denoted as sgCon); sgRNAs targeting *Cdk1* or *Pten* were used as positive controls that were expected to produce cancer blocking or promoting effects, respectively ^30,31^. Initial analysis using liver to body weight ratios (LW/BW) showed that for 7 of 11 genes tested, the associated sgRNAs trended to reduced tumor burden, supporting the observation that *CTNNB1-*mutant tumors arise more readily in zone 3 (**Fig. 4b**). The mean LW/BW of sgCdk1 mice (n=11) and sgPten mice (n=5) were smaller and larger than sgCon mice, respectively. Reduction of targeted mRNAs and proteins was observed in the normal liver as well as in isolated liver tumors (**Fig. 4c, Extended Data Fig. 9a-c**).

## Gstm2 or Gstm3 loss inhibits CTNNB1/MYC induced HCC initiation

Deletion of either *Gstm2* or *Gstm3*, which are zone 3 specific genes, led to significantly reduced LW/BWs compared to the sgCon group (**Fig. 4d**). Gstm’s are Glutathione transferase family members within the mu sub-family that are thought to be activated upon WNT signaling ^32^. In human HCCs, higher expression of *Gstm3* is associated with shorter survival and worse prognosis (**Extended Data Fig. 9d**) ^33,34^. To examine overall survival in a less cancer-prone strain background, we repeated these experiments in C57BL/6J mice and found the same pattern, with *Gstm3* or *Gstm2* deletion reducing LW/BW ratios (**Extended Data Fig. 9e,f**) and improving survival (**Fig. 4e**). SgGstm3 and sgGstm2 KO livers showed a 5-fold decrease in the relative area of microscopic tumors (**Fig. 4f,g**). We tracked mutant cells with GS as a marker. Three weeks after HDT, there was a large increase in liver tumors and in GS+ staining in sgCon vs. sgGstm3 livers (**Fig. 4h**). To determine if increased *Gstm3* could promote tumors, we overexpressed *Gstm3* together with *CTNNB1* and *MYC*, which trended to an increase in tumor burden 1 week after HDT (**Extended Data Fig. 9g**). To determine if Gstm’s are also required for cancer cell maintenance, we generated *Gstm3* KO HepG2 cells (**Extended Data Fig. 9h)**. These KO cells showed a significant reduction in proliferation (**Extended Data Fig. 9i**). Taken together, we found that zone 3-specific Gstm’s were necessary and sufficient to accelerate *CTNNB1-*driven liver carcinogenesis.

The importance of *Gstm3* in promoting HCC could be related to its normal metabolic function in zone 3 hepatocytes. GSTM3 plays an important role in xenobiotic detoxification and ROS scavenging near the CV, where metabolites and toxins exit the liver. Detoxification of xenobiotics and ROS is thought to play a key role in both anticancer drug resistance and tumorigenesis ^35,36^. *Gstm3* deletion led to an increase in apoptotic hepatocytes compared to control livers (**Fig. 4i, Extended Data Fig. 9j**). Concurrently, peroxide quantification and superoxide staining using DHE showed an increase in ROS levels in *Gstm3* KO vs. control livers, indicating that the loss of *Gstm3* and its antioxidant activity leads to ROS accumulation (**Fig. 4j, Extended Data Fig. 9k**). Thus, *Gstm3* likely promotes HCC initiation through pro-survival, anti-oxidant functions within mutant zone 3 hepatocytes that would otherwise be outcompeted by WT hepatocytes.

## Therapeutic inhibition of GSTM2/3 can prevent liver cancer initiation

To ask if the knowledge of zone-specific origins of HCC could help identify preventative interventions, we tested a chemical inhibitor of GSTM activity. Ethacrynic acid is a loop diuretic that has been found to inhibit GST-family proteins with more potency against the GSTM class, in part by acting as an alternative substrate for glutathione conjugation ^37,38^. Mice were randomized to receive vehicle control or ethacrynic acid treatment starting two days prior to HDT, and then all mice were hydrodynamically transfected with *CTNNB1* and *MYC* plasmids (**Fig. 4k**). There was reduced LW/BW in the ethacrynic acid group compared to the vehicle control PBS group (**Fig. 4l**), as well as a 4-fold reduction in microtumor area (**Fig. 4m,n**). Similarly, treatment with ethacrynic acid led to increased peroxide levels compared to vehicle treatment, indicating successful inhibition of GST activity (**Fig. 4o**). Therefore, chemical inhibition of GSTM proteins recapitulates the tumor inhibitory effects of *Gstm2* and *Gstm3* deletion.

## DISCUSSION

The observation that clonal hematopoiesis is a high risk condition for leukemogenesis has supported the paradigm that stepwise expansion of premalignant mutant clones is fundamental to leukemia evolution ^39,40^. Recent sequencing studies, including those from non-malignant liver tissues have revealed that somatic mutations are very common, but mutations in cancer drivers are observed less frequently than anticipated ^9–11^. This suggests that clones containing mutations with the potential to induce malignancy do not necessarily progress to cancer through stepwise clonal expansion. To understand the fates of cells that undergo the first steps in malignant transformation, we modeled these truncal mutations in normal cells and visually followed mutant clone fates on the path toward carcinogenesis. We needed to overcome a major challenge in studying clonal evolution in solid tissues, which is that normal cells with mutations are not easily traceable. Cre-lox technologies cannot reliably induce both a mutation and reporter activation in the same clones. The case of *Ctnnb1* in the liver is unique, and allowed for unprecedented resolution in mutant clone lineage tracing. Using expression of GS, we successfully lineage-traced mutant hepatocytes prior to carcinogenesis, affording us the ability to quantify *Ctnnb1* mutation burden over space and time.

Perhaps the most unexpected result was that mutant clones in different locations in the liver lobule have dramatically divergent fates. Mutant clones adjacent to portal tracts expanded to maintain the GS+ population in zone 1, while mutant clones in zones 2 and 3 disappeared. We believe that the basis for this observation is the differences in the relative competitiveness between WT and mutant clones in distinct zones. In zone 3, most WT hepatocytes are already exposed to WNT signals and are thus more competitive against *Ctnnb1* mutant clones, while in zone 1, WT hepatocytes are WNT-inactive and less proliferative than mutant clones. While we invoke a simplistic model involving cell autonomous fitness differences, zone specific immunosurveillance and antagonistic cell-cell signaling are also possible.

Not only was it surprising that zonation correlated with drastically different fates, we found that zone 3 mutants developed HCC much more frequently than zone 1 mutants. This demonstrated that the degree of clonal expansion of mutant hepatocytes did not at all predict their tumorigenic potential. Unlike what occurs in clonal hematopoiesis, our observations in the liver suggest that clonal expansion is not essential for tumorigenesis, and in the case of *Ctnnb1*, is anti-correlated with cancer development. In zone 3, there is a well-known histologic pattern of liver damage in diseases that culminate in HCC ^41,42^, and zone 3 is frequently associated with oxidative stress ^43,44^. It stands to reason that antioxidant enzymes are needed to protect wild-type zone 3 cells from dying in a harsh environment. As a consequence of oxidative stress, mutations may accumulate more frequently in cells within zone 3. It is likely that WT cell competition against mutant clones within a cirrhotic nodule is a key tumor suppressive mechanism. However, the existence of many antioxidant mechanisms in this zone, including GST related enzymes, become co-opted for tumor cell survival. Ultimately, cell competition is not able to suppress transformation when zone 3 cells survive to accumulate the requisite mutations, in part through antioxidant mechanisms that are native to zone 3.

One genetic contributor to the increased rate of transformation appears to be the zonated expression of enzymes such as GSTM2/3, antioxidant phase II detoxification enzymes that perform glutathione conjugation of xenobiotics. Previously, it was shown that GSTM2 binds to and inhibits the MAPK effector kinase ASK1, which is known to drive apoptosis upon stress signaling in the liver. ASK1 deletion in the liver also promoted cancer formation ^45^. Altogether, it is likely that GSTM proteins mediate multiple effects including antioxidant and anti-apoptotic ones, which support transformation within mutant clones in zone 3. Furthermore, we also identified several other zone 3 genes that promote tumorigenesis (**Fig. 4b**), indicating that Gstm’s are not alone in their pro-tumor functions. Given that many zone 3 metabolic enzymes can be modulated with small molecules, our findings suggest strategies for cancer prevention that capitalize on knowledge of zonation dependent mechanisms of liver carcinogenesis.

## Methods

### Animals

All mice were handled in accordance with and with the approval of the Institutional Animal Care and Use Committee (IACUC) at UT Southwestern. *ApoC4-CreERT2*; *Rosa26^LSL-tdTomato/+^, Gls2-CreERT2*; *Rosa26^LSL-tdTomato/+^,* and *Cyp1a2-CreERT2*; *Rosa26^LSL-tdTomato/+^* mice were generated by Dr. Yonglong Wei ^19^ on C57/BL6 strain background. *Arid2^fl/fl^* mice ^46,47^ were obtained from the Mutant Mouse Resource & Research Center (MMRRC Program). *Ctnnb1^(ex3)^ ^fl/fl^* mice ^48^ on a Swiss Webster strain background were obtained from the UTSW O’Brien Kidney Research Core. First, *CreERT2; Rosa26^LSL-tdTomato/+^* mice were crossed to *Arid2^fl/fl^* mice. Progeny male *CreERT2; Arid2^fl/+^;Rosa26^LSL-tdTomato/+^*mice were then crossed to *Arid2^fl/fl^* mice to obtain male *CreERT2; Arid2^fl/fl^;Rosa26^LSL-tdTomato/+^* mice. Separately, *Ctnnb1^(ex3)^ ^fl/fl^* mice were crossed to *Arid2^fl/fl^* mice. Progeny male *Ctnnb1^(ex3)^ ^fl/+^; Arid2^fl/+^* mice were crossed to female *Arid2^fl/fl^* mice to obtain female *Ctnnb1^(ex3)^ ^fl/+^; Arid2^fl/fl^* mice. Finally, male *CreERT2; Arid2^fl/fl^;Rosa26^LSL-tdTomato/+^* mice were crossed to female *Ctnnb1^(ex3)^ ^fl/+^; Arid2^fl/fl^* mice to generate experimental animals that were *CreERT2; Ctnnb1^(ex3)^ ^fl/+^; Arid2^fl/fl^;Rosa26^LSL-tdTomato/+^*or *CreERT2; Ctnnb1^(ex3)^ ^fl/+^; Arid2^fl/fl^*. This cross also generated *CreERT2; Arid2^fl/fl^;Rosa26^LSL-tdTomato/+^* or *CreERT2; Arid2^fl/fl^* mice without the *Ctnnb1^(ex3)^ ^fl/+^* activation allele, as well as Cre-negative *Ctnnb1^(ex3)^ ^fl/+^; Arid2^fl/fl^* and *Arid2^fl/fl^* control mice. Notably, the cross above does not generate animals with only the *Ctnnb1* mutant activation allele. In order to generate those mice, we had to use an independent crossing strategy. *CreERT2; Rosa26^LSL-tdTomato/+^*mice were crossed to mixed background *Ctnnb1^(ex3)^ ^fl/+^; Rosa26^LSL-tdTomato/+^* to generate *CreERT2; Ctnnb1^(ex3)^ ^fl/+^; Rosa26^LSL-tdTomato/+^* mice. Genotyping primers are available in **Extended Data Table 7**. For all *CreERT2* genotyping, both L and R primer pairs are used to confirm the correct CreER strain.

At 4 weeks of age, experimental animals were weaned from their mothers and injected with 100 mg/kg tamoxifen for 2 consecutive days on postnatal days 28 and 29. 100 mg of tamoxifen (Sigma-Aldrich #T5648-1G) was mixed by vortexing in 10 mL of corn oil (Sigma-Aldrich #C8267-500ML) and sonicated in a water bath for up to 30-45 minutes, with vortexing every 10 minutes to help the tamoxifen go into solution. Once the tamoxifen was fully dissolved in corn oil, the solution was aliquoted and frozen at -20°C for storage. Four week old mice were weighed and injected intraperitoneally (IP) with the tamoxifen 10 mg/mL solution with a volume (μL) 10 times their weight in grams. This resulted in a dose of 100 mg/kg tamoxifen.

### TCGA mutation analysis

To determine the likelihood of various driver genes being co-mutated with *CTNNB1* in human HCC, we used the cBio Cancer Genomics Portal ^49,50^. We queried previously published HCC data sets ^2,51–53^ for the following genes: *TERT, TP53, CTNNB1, ALB, APOB, AXIN1, ARID1A, ARID2*, *NCOR1, RB1, CDKN2A, ERRFI1, PTEN, CCND1*. Then we looked under the “Mutual Exclusivity” tab and examined combinations that include *CTNNB1*. We specifically examined the log_2_ Odds Ratio, which “quantifies how strongly the presence or absence of alterations in A are associated with the presence or absence of alterations in B in the selected samples.”

### Immunostaining and histology

For immunofluorescence, mice were sacrificed and livers were collected in 4% paraformaldehyde (Thermo Fisher Scientific #AAJ19943K2) and shaken overnight at 4°C. Then the livers were transferred into a sucrose (30% w/v) in PBS (Thermo Fisher Scientific #SH3002802) solution and shaken overnight at 4°C. Chunks of liver were then cryoembedded in Tissue-Tek® O.C.T. Compound (Sakura Finetek USA inc, VWR #25608-930) and sectioned using a cryostat. 8-9 μm liver sections were mounted onto glass microscope slides. Liver sections were then washed with PBS for 2 min x2 to remove residual O.C.T. compound. Liver sections were blocked in 5% bovine serum albumin (BSA, Sigma Aldrich #A3294-500G) + 0.25% Triton X-100 (Sigma Aldrich #X100-100ML) dissolved in PBS for 1 hour at room temperature. Then, primary antibodies were diluted in blocking solution. For glutamine synthetase (GS) staining, primary antibody (Abcam #ab49873) was dissolved 1:1000 in blocking solution. For Ki-67 staining, primary antibody (Abcam #ab15580) was dissolved 1:500 in blocking solution. Liver sections were incubated in primary antibody solution overnight at 4°C. The next day, sections were washed in PBST (0.1% Tween 20, Thermo Fisher Scientific #BP337500) for 5 min, 3X. Then secondary antibody (Goat anti-Rabbit IgG (H+L) Cross-Adsorbed Secondary Antibody, Alexa Fluor 488, ThermoFisher #A-11008) was diluted 1:100 in blocking solution. Liver sections were incubated in secondary antibody for 1 hour at room temperature. Then sections were washed in PBST for 5 min, 3X. Liver sections were mounted with VECTASHIELD® PLUS Antifade Mounting Medium with DAPI (Vector Laboratories #H-2000-10) and coverslipped. Stained and mounted slides were imaged using a standard fluorescent microscope system. Whole section immunofluorescence images were obtained using a Zeiss Axioscan.Z1 digital slide scanner through the UT Southwestern Whole Brain Microscopy Facility.

For immunohistochemistry, tissue was paraffin embedded and sectioned by the UTSW Histopathology core. Sections were deparaffinized twice with xylene and rehydrated with a graded series of ethanol (100% twice, 90%, 80%, 50%, 30% ethanol, and distilled water) for 5 min each. Antigen retrieval was performed for 20 min in Antigen Retrieval Citra Plus buffer (BioGenex #HK080) at sub-boiling temperature and allowed to come to room temperature for 30 min. Sections were washed in distilled water and incubated for 10 min in 3% (vol/vol) hydrogen peroxide in methanol to block endogenous peroxidase activity. Sections were then washed in PBST for 5 min and blocked at room temperature with M.O.M. Mouse Ig Blocking Reagent (Vector Laboratories #PK2200) for 1 hour at room temperature. Next, sections were washed in PBST for 2 min, 2X and incubated with M.O.M. Protein Concentrate Solution (Vector Laboratories #PK2200) in PBST for 5 min at room temperature. Primary antibodies were diluted in 10% goat serum in PBST. For beta-catenin staining, primary antibody (BD Transduction Laboratories #610154) was dissolved 1:500 in goat serum. Liver sections were incubated in primary antibody solution overnight at 4°C. The next day, sections were washed in PBS for 2 min, 2X, and incubated in Biotinylated Anti-Mouse IgG Reagent (Vector Laboratories #PK2200) for 10 min at room temperature. After washing in PBS for 2 min, 2X, sections were incubated with avidin/biotin conjugation solution (Vector Laboratories #PK6100) for 30 minutes at room temperature. Sections were washed in PBS for 5 min, 2X, and then developed using a DAB substrate kit (Vector Laboratories #SK4100). Sections were finally counterstained with hematoxylin (Vector Laboratories #H4304), mounted with Aqua-Mount (Epredia #13800), and coverslipped.

For H&E staining, livers were collected after sacrificing in 4% paraformaldehyde as described above and then incubated in 70% ethanol. Tissue was paraffin embedded, sectioned, and stained by the UTSW Histopathology Core. Whole section H&E and IHC images were obtained using a Hamamatsu NanoZoomer 2.0-HT digital slide scanner through the UT Southwestern Whole Brain Microscopy Facility.

### 5-ethynyl-2’-deoxyuridine proliferation assay

1 month after tamoxifen, mice were provided with autoclaved water containing 1 mg/mL 5-ethynyl-2’-deoxyuridine (EdU, Carbosynth #NE08701) for 10 days. After 10 days, mice were sacrificed and livers were collected and cryosectioned as described above for immunofluorescence. EdU signal was detected in frozen sections using the Click-iT EdU Alexa Fluor 488 Imaging Kit (Life Technologies #C10337) following the manufacturer’s instructions. Briefly, sections were further fixed in 4% paraformaldehyde for 15 min at room temperature and washed in 3% BSA dissolved in PBS for 2 min, 2X. Sections were then permeabilized in 0.5% Triton X-100 in PBS for 20 min at room temperature and washed in 3% BSA dissolved in PBS for 2 min, 2X. The Click-iT reaction cocktail was prepared, and sections were incubated overnight at 4°C. The next day, sections were washed in 3% BSA dissolved in PBS for 2 min, 1X, blocked, and stained for GS as described above. Stained and mounted slides were imaged using a standard fluorescent microscope system.

### Single nuclear RNA sequencing

Primary mouse hepatocytes were isolated using a previously described two-step collagenase perfusion method ^54^. The perfused hepatocytes were gently washed and resuspended in 25 mL Hepatocyte (1X) Wash Buffer (Gibco #17704024) and centrifuged at 50g for 5 minutes at 4°C The supernatant was discarded and the wash step was repeated two additional times. After the two wash steps, the hepatocytes were resuspended in 25 mL Hepatocyte (1X) Wash Buffer. Using Trypan Blue dye, the cell number was counted. 1 X 10^6^ cells were transferred to a 1.5 mL microtube and centrifuged at 500g for 5 minutes at 4°C. The supernatant was discarded and hepatocytes were resuspended in 1 mL GBSS (Sigma #G9779) buffer supplemented with 0.7% CHAPS (Thermo #28299) and 0.2U/μl RNase inhibitor (Roche #3335399001). Hepatocytes were lysed on ice for 5 minutes. Then they were centrifuged at 500g for 5 minutes and the isolated nuclei were resuspended in nuclear wash buffer: 1 mL PBS buffer supplemented with 1.0% bovine serum albumin (BSA) and 0.2U/μl RNase Inhibitor. Wash was repeated in the nuclear wash buffer once. The nuclei were resuspended in 200 μl nuclear wash buffer and nuclei number were counted using Trypan Blue dye. Single nuclei libraries were prepared using the 10x Genomics Chromium Next GEM Single Cell 5′ Library and Gel Bead Kit v1.1 (10X Genomics #1000167). Whole transcriptome libraries were generated according to the manufacturer’s protocol. For each sample, about 8,000 nuclei were mixed with reverse transcription master mix and loaded to Next Gem Chip-G to target ∼5,000 nuclei after recovery. Libraries were sequenced using 150 bp paired-end Illumina NextSeq500 system at the UTSW Children’s Research Institute Sequencing Facility.

### Single nuclear RNA sequencing analysis

10x scRNA-seq data was pre-processed using the Cell Ranger software (4.0.0) provided by 10X Genomics. We used the “mkfastq”, “count” and ‘aggr’ commands to process the 10x scRNA-seq output into one cell by gene expression count matrix, using default parameters. A custom-built version of mouse genome assembly GRCm38 which included unspliced pre-mRNA was used. (https://support.10xgenomics.com/single-cell-gene-expression/software/pipelines/latest/advanced/referen ces) scRNA-seq data analysis was performed with the Scanpy (1.6.0, ref 2) package in Python ^55^. Genes expressed in fewer than 3 cells were removed from further analysis. Cells expressing less than 100 and more than 6000 genes were also removed from further analysis. In addition, cells with a high (>= 0.1) mitochondrial genome transcript ratio were removed. For downstream analysis, we used count per million normalization (CPM) to control for library size differences in cells and transformed those into log(CPM+1) values. After normalization, we used the ‘pp.highly_variable_genes’ command in Scanpy to find highly variable genes across all cells using default parameters except for “min_mean = 0.0125 and min_disp=0.5”. The data were then z-score normalized for each gene across all cells. We then used the ‘tl.pca (default parameters)’, the ‘pp.neighbors (n_neighbors=25)’ and the ‘tl.leiden (resolution = 0.2)’ commands in Scanpy to partition the single cells into distinct clusters. Briefly, these processes first identify 50 principle components in data based on the previously found highly variable genes to reduce the dimensions in the original data, and then build a nearest neighbor graph based on the top 25 principle components, and finally a partition of the graph that maximizes modularity was found with the Leiden algorithm ^56^. The most highly expressed genes from each cluster were found through contrasting a cluster of cells with all the other cells (self-vs-rest) using the Wilcoxon signed-rank test, which is included in the “tl.rank_genes_groups” function in the Scanpy package. We predicted the cell cycle phase by scoring each single cell against a reference cell cycle gene list ^57^ using the “tl.score_genes” function in the Scanpy package. Briefly, the cell cycle score was calculated as the average expression of the target gene set subtracted with the average of a random set of reference genes.

### Computational image analysis to quantify GS

We segmented two areas from each immunofluorescent image in our analysis. The tissue masks were defined as areas with no major blood vessels, whose intensity values were similar to image background on DAPI channel. We used the Otsu thresholding [M. Sezgin & B. Sankur (2004). "Survey over image thresholding techniques and quantitative performance evaluation". Journal of Electronic Imaging. 13 (1): 146–165. doi:10.1117/1.1631315] to estimate the background intensity and the final DAPI background was set to 0.2-fold of the Otsu threshold. The GS masks were defined as areas with positive GS stain, which were also determined using the Otsu thresholding. All images were preprocessed with adaptive histogram equalization and all tissue masks were smoothed with a Gaussian filter with sigma = 2. Using this method, we were able to quantify the number of pixels in the GS mask and the tissue mask (Extended Data Fig. 1). We divided the number of pixels in the GS mask by the number of pixels in the tissue mask and multiplied this ratio by 100, to get a percentage of total tissue area that is GS+ in each image.

### Quantification of zonal position index

We used a previously described measure called the position index (P.I.), determined by the distance to the closest central vein (CV) (x), the distance to the closest portal triad (PT) (y), and the distance between the CV and PT (z), based on the law of cosines ^26^: 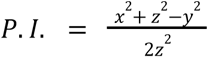 The P.I. is a value between 0 and 1 that reflects a hepatocyte’s relative position along the CV - PT axis. We subdivided these values and this axis into 9 layers such that layers 1-3 are pericentral zone 3, layers 4-6 are midlobular zone 2, and layers 7-9 are periportal zone 1. After installing a publicly available ImageJ macro called “Measure and Label” (https://imagej.nih.gov/ij/plugins/measure-label.html), we used our GS stained images to measure the x, y, and z distances, and calculated P.I. for mutant hepatocytes in mutant mice.

### Digital droplet PCR (ddPCR)

A primer / probe set (HEX fluorophore) was designed for the *Ctnnb1* floxed allele and another primer probe set (FAM fluorophore) was designed for the *Ctnnb1* exon 3 deletion allele (primer/probe sequences are available in **Extended Data Table 8**). The amplicon for the *Ctnnb1* floxed allele is entirely contained within the floxed region. The amplicon for the *Ctnnb1* deleted allele (207 bp) can easily be amplified from an exon 3 deleted allele but the same primer set spans a very large (2.4 kb) region in a floxed DNA allele, which would not be amplified (**Extended Data Fig. 2**). The droplet reader measures the number of HEX+ and/or FAM+ droplets, and the ddPCR assay output for each sample is FAM copies / µL and HEX copies / µL. By dividing (FAM copies / µL) / [(FAM copies / µL) + (HEX copies / µL)] X 100, we calculated the percentage of *Ctnnb1* exon 3 deleted alleles for each sample. Then we compared the percentage of *Ctnnb1* mutant alleles relative to the initial time point of 1 month after tamoxifen (100% relative exon 3 deletion). Genomic DNA samples were extracted from liver tissue using the PureLink™ Genomic DNA Mini Kit (Invitrogen #K1820-01) per the manufacturer’s protocol for tissue samples. For ddPCR, samples were prepared according to BioRad’s guidelines. In brief, custom ddPCR primer/probes were designed as described above and ordered from BioRad. Then, ddPCR samples were prepared with gDNA from liver samples, primer/probes, and ddPCR Supermix for Probes (No dUTP) (Biorad #1863024). Droplets were generated using the QX200 Droplet Digital PCR System per the manufacturer’s protocols using Droplet Generation Oil for Probes (BioRad #1863005) and DG8 Cartridges and Gaskets (BioRad #1864007). PCR was performed with the following cycle: 1. Ramp (2°C/s) to 95°C, for 10 min.; 2. Ramp (2°C/s) to 94°C for 30 sec; 3. Ramp (2°C/s) to 55°C for 1 min.; 4. Go to step 2 40X; 5. Ramp (2°C/s) to 98°C for 10 min.; 6. Hold at 4°C. PCR plates were left at 4°C overnight. The following day, droplets were read with the QX200 Droplet Digital PCR System per the manufacturer’s protocols after sample names and appropriate fluorophores were input into the template.

### Tumor quantification

At 6-12 months post-tamoxifen injection, animals were sacrificed and livers were collected. We evaluated the livers for tumor burden by taking photos of each side of the livers and then counting surface tumor number as a surrogate for overall tumor burden. After photos were taken, tumors were dissected to avoid any non-tumor tissue and placed into 1.5 mL tubes. These tubes were then snap frozen in liquid nitrogen for downstream assays and stored at -80°C.

### RNA sequencing

Fragments of frozen tumors were homogenized in TRIzol (TRIzol Plus RNA Purification Kit, Life Technologies #12183555) using a tissue homogenizer. Then, RNA was extracted using the PureLink™ RNA Mini Kit (Invitrogen #12183018A) according to the manufacturer’s protocol (page 50, “Using TRIzol® Reagent with the PureLink® RNA Mini Kit”). Additionally, on-column DNAse treatment was performed with PureLink™ DNase Set (Invitrogen #12185010) according to the manufacturer’s protocol (page 63, “On-column PureLink® DNase Treatment Protocol) prior to elution of RNA. Libraries were prepared with the Ovation RNA-Seq Systems 1-16 (Nugen) using 100 ng of DNAse-treated RNA and indexed libraries were multiplexed in a single flow cell and underwent 75 base pair single-end sequencing on an Illumina NextSeq500 using the High Output kit v2 (75 cycles) at the Children’s Research Institute at UTSW Sequencing Core Facility. Sequencing reads were aligned with hisat2 (version 2.1.0) with option –rna-strandness F to mm10. Gene count reads were counted with HTseq (version 0.6.1). Differentially expressed genes were called with DESeq (version 1.11.4) with default settings.

### Real-time quantitative PCR

RNA was isolated from tumors as described above and RNA concentration was determined. Reverse transcription was performed on 1 μg of RNA using the iScript™ cDNA Synthesis Kit (Bio-Rad #1708891). Then, real-time qPCR was performed on a Bio-Rad CFX96 Touch Deep Well Real-Time PCR System using iTaq™ Universal SYBR® Green Supermix (Bio-Rad #1725125) according to the manufacturer’s protocol. qPCR primer sequences were obtained from https://pga.mgh.harvard.edu/primerbank/. Sequences are listed in **Extended Data Table 9**.

### Identification of zonated genes for *in vivo* and *in vitro* CRISPR screening

We used bulk RNA sequencing data published by Ben-Moshe et al ^17^. Briefly, their work used expression of two zonated cell surface proteins, CD73 and E-cadherin, to sort 8 different hepatocyte populations. Bulk RNA sequencing showed that the gene expression of these 8 spatially sorted hepatocyte populations was highly concordant with the zonated expression profile from their previous single-cell RNA sequencing experiments ^27^. For our purposes, we wanted to select the genes that were most highly specific to either periportal or pericentral hepatocytes.

First, we calculated the average expression of each gene (given as unique molecular identifier counts per million in the Extended Table 2 from their paper) in each layer across 5 unique mice, where layer 1 is most pericentral and layer 8 is most periportal. Then, we calculated the maximum expression of each gene across the 8 layers, and eliminated all genes where the maximum average expression was < 50 counts per million (CPM), assuming that these are genes that are lowly expressed throughout the liver. Then, we calculated the linear R^2^ value for each gene across the 8 layers and eliminated all genes with an R^2^ < 0.75, thus assuming that genes that peak in periportal or pericentral hepatocytes exhibit more linear expression across layers. Then, we calculated the slope of the linear regression line for each gene, assuming that more zonated genes would exhibit a steeper slope. Genes with a slope > 0 are more periportal while genes with a slope < 0 are more pericentral. However, given that some genes are very highly expressed across all layers (such as *Alb*), the slope was divided by the average expression of that gene across all layers. This gave us a “normalized slope.” Finally, R^2^ was multiplied by “normalized slope” for each gene to give a “zonation score.” Genes were sorted by zonation score. Those with the highest, or most positive, zonation score are highly zonated periportal genes, while those with the lowest, or most negative, zonation score are highly zonated pericentral genes. The 80 most periportal and 80 most pericentral genes were selected to be included in our CRISPR screens.

Independent of the first selection method, we additionally used a second method to select genes. To select for genes most highly expressed in periportal hepatocytes, genes with expression < 50 CPM in layer 8 or > 300 CPM in layer 1 were eliminated. Then, layer 8 - layer 1 expression (difference) was calculated and layer 8 / layer 1 expression (ratio) was calculated. The difference was multiplied by the ratio to give a “periportal score.” Genes were ranked and sorted by periportal score. The top 80 genes were selected as highly zonated periportal genes. To select for highly zonated pericentral genes, the same calculations were performed, but in the inverse. Genes with expression < 50 CPM in layer 1 or > 300 CPM in layer 8 were eliminated. The difference of each gene between layer 1 and layer 8 and the ratio of layer 1 to layer 8 were calculated. The difference was multiplied by the ratio, and genes were ranked and sorted by this “pericentral score.” The top 80 genes were selected as highly zonated pericentral genes. These two methods of gene selection for the screen found 48 common periportal genes and 60 common pericentral genes. The lists were combined, resulting in a final list of 111 periportal genes and 103 pericentral genes (**Extended Data Table 6**).

### *In vivo* CRISPR screening

We designed two libraries against our target genes, one for the CRISPR knockout (CRISPR KO) screen and another for the CRISPR activation (CRISPRa) screen. sgRNA sequences for the CRISPR KO screen were extracted from the mouse Gecko v2 library ^58^, which were designed against coding exons. Notably, 23 genes that we had selected were not found in the Gecko v2 library, so these were excluded. The final CRISPR KO library consisted of 1170 sgRNAs, which included 6 guides targeting each of the 191 genes and 23 non-targeting control guides. The sgRNAs were synthesized and subcloned into our transposon vector with good representation (> 99%) and high uniformity across sgRNAs.

sgRNA sequences for the CRISPRa screen were extracted from the mouse Genome-wide CRISPRa-v2 library ^59^, which target sites upstream of the transcriptional start site (TSS) of a given gene. Notably, 26 genes that we had selected were not found in the Genome-wide CRISPRa-v2 library, so these were excluded. The final CRISPRa library consisted of 1026 sgRNAs, which included 5 guides targeting each of the 188 genes and 21 non-targeting control guides. The sgRNAs were synthesized and subcloned into our transposon vector with good representation (> 99%) and high uniformity across sgRNAs.

For both screens, 5 mg of plasmid and 1 mg of SB100 transposase plasmid were delivered via HDT into *Fah* KO mice (n = 10 for KO screen, n = 7 for activation screen). NTBC water was withdrawn immediately after injection. After 1 month, the liver was collected and DNA was extracted from the whole liver. The sgRNA was amplified and sequenced on a NextSeq500. sgRNA representation was analyzed by the MAGeCK algorithm using the default settings.

### Hydrodynamic transfection (HDT)

SB100, pT3-Myc, and Ctnnb1 transposon plasmids were obtained from Xin Chen at UCSF. We selected 11 highly enriched zone 3 genes for CRISPR-mediated knockout from our sequencing and screening results. For each gene, pairs of sgRNAs were generated using the CRISPick tool from the Broad Institute ^60,61^ and sequentially cloned into vector px333 (Addgene, #64073, Cambridge, MA, USA). px333 contains two sgRNA cassettes with unique digestion sites for BbsI and BsaI. GFP, Luciferase, and Gstm3 expression transposon plasmids were modified from pT3-EF1a-GW, which was obtained from Xin Chen at UCSF. cDNA for GFP and Luciferase were synthesized by *GENEWIZ* and cDNA for Gstm3 were synthesized from Genscript. Gateway sites were added to the cDNA through PCR and genes of interest were then used to replace the CcdB gene by Gateway recombination (Invitrogen #11789100 and #11791100). Male FVB or C57BL/6J mice were injected at approximately 6 weeks of age when their body weight was at least 20 g. HDT plasmids were suspended in 2 mL of saline and administered via tail vein injection in 7 seconds. A 10:1 mass ratio of combined HDT plasmids to transposase plasmid was used (e.g. 10 μg pT3-Ctnnb1-ΔN90 + 10 μg pT3-cMyc + 10μg tandem zone 3 sgRNA px333 : 3 μg SB100 transposase).

### Ethacrynic acid experiments

For chemical inhibition of GST activity, experimental animals were intraperitoneally injected with 10 mg/kg ethacrynic acid (Santa Cruz Biotechnology #SC-257424) starting two days before HDT injection. Mice were given ethacrynic acid daily thereafter until sacrifice. 1 g of ethacrynic acid was initially dissolved 50 mL of 100% ethanol. This stock solution was then diluted 20x in DPBS (Sigma Aldrich #D5652) to a final concentration of 1 mg/mL for use. The 7-week old mice were weighted and injected with the ethacrynic acid 1 mg/mL solution with a volume (μL) 10 times their weight in grams.

### TUNEL Staining

To assess apoptosis in tissue sections, TUNEL staining with the In Situ Cell Death Detection Kit, Fluorescin (Roche #11684795910) was used to detect double stranded DNA breaks according to the manufacturer’s instructions. Briefly, paraffin-embedded tissue sections were deparaffinized in xylenes and rehydrated with a graded series of ethanol as described above. Then, sections were incubated with Proteinase K (Invitrogen #25530049) diluted 1:1000 in PBS for 30 min at 37°C in a humidified chamber. After washing in PBS 2 min, 2X, the sections were incubated in TUNEL reaction mixture for 1 hour at 37°C in a humidified chamber. Finally, sections were washed in PBS 2 min, 2X, mounted with VECTASHIELD® PLUS Antifade Mounting Medium with DAPI (Vector Laboratories #H-2000-10) and coverslipped. Stained and mounted slides were imaged using a standard fluorescent microscope system

### ROS detection

To quantify reactive oxygen species (ROS), the ROS-Glo H_2_O_2_ Assay Kit (Promega #G8820) was used according to the manufacturer’s instructions. Briefly, 20 mg of tissue was lysed with Passive Lysis 5X Buffer (Promega #E1941), and the lysate was incubated with H_2_O_2_ Substrate Solution at room temperature for 1 hour. Afterwards, ROS-Glo Detection Solution was added for 20 minutes at room temperature, and luminescence was measured with a FLUOstar Omega Microplate Reader. To detect ROS, mice were sacrificed and livers were either snap-frozen and stored in -80°C or directly cryoembedded and cryosectioned as described previously. 10 μm sections were mounted onto glass microscope slides and incubated with 10 μM dihydroethidium (DHE, Thermo Fisher Scientific #50850563) at 37°C for 30 min as previously described ^62^. The resulting red fluorescence intensity corresponds to cellular superoxide anion levels.

### Western blot analysis

HDT livers were collected and homogenized in T-PER Tissue Protein Extraction Reagent (Life Technologies #78510) supplemented with protease (ApexBio #K1007) and phosphatase (ApexBio #K1012) inhibitor cocktails according to the manufacturer’s instructions. Cells were spun down and lysed in RIPA buffer (Thermo Fisher Scientific #PI89900) supplemented with protease and phosphatase inhibitor tablets (Life Technologies #A32961). Lysates were centrifuged at 8,000 g for 10 min and the supernatants were collected and stored at -80C. The following antibodies were used: anti-GSTM3 (Abcam ab272613 or ab229858), anti-β-Tubulin (CST #2128), anti-rabbit IgG, HRP-linked Antibody (CST #7074), anti-mouse IgG, HRP-linked Antibody (CST #7076).

### Cell culture

HEK293T and HepG2 cells were cultured in Dulbecco’s modified Eagle’s medium (DMEM) with 10% fetal bovine serum (FBS) and 10% penicillin-streptomycin. SNU398 cells were cultured in RPMI medium with 10% heat-inactivated FBS and 10% penicillin-streptomycin. All cells were cultured at 37 °C in a 5% CO2 incubator.

### Lentiviral particle production

*Gstm3* sgRNA oligos were inserted into the lentiCRISPRv2 backbone plasmid through digestion of vectors with BsmbI. For virus production, 3.5 μg of the appropriate plasmid and 3.2 μg of helper plasmids (2.7 μg of gag-pol and 0.5 μg of VSVg) were transfected into HEK293T cells cultured at 50% confluence in a 10 cm dish using Lipofectamine 3000 (Invitrogen) according to the manufacturer’s instructions. Viral supernatants were collected 48 hr after transfection and filtered through a 0.45 μm filter.

### Statistics

Statistical tests used are noted in the figure legends. Unless otherwise stated in the methods or figure legends, two-tailed Student’s t tests (two-sample equal variance) were used to test the significance of differences between two groups. Variation is indicated using standard error of the mean (SEM) and presented as mean ± SEM. Statistical significance is displayed as * (p < 0.05), ** (p < 0.01), *** (p < 0.001), and **** (p < 0.0001).

## Acknowledgements

We thank Shawn Burgess, Ralph Deberardinis, and David Hsieh for constructive comments on the manuscript; C. Lewis (UTSW Tissue Procurement Service) and J. Shelton (UTSW Histopathology Core) for histopathology; D. Ramirez (UTSW Whole Brain Microscopy Facility) for imaging; J. Xu and Y. J. Kim (CRI Sequencing Core) for sequencing. H.Z. is supported by the Pollack Foundation, NIH R01 grants (CA251928, AA028791, DK125396), and the Emerging Leader Award from the Mark Foundation For Cancer Research (#21-003-ELA). The Zhu lab and J.G. are supported by an Innovation Award from the Moody Medical Research Institute.

## Author contributions

A.C., J.G. and H.Z. conceived the project, performed the experiments, and wrote the manuscript. performed animal and cell culture experiments. P.G. performed the histologic analysis. Y.J., Y.W., G.X., and T.W., performed bioinformatic analysis of images and sequencing data. N.C., L.L., Y.W., M.Z., Z.W., and H.G. assisted with animal and cell culture experiments.

## Conflicts statement

H.Z. has a sponsored research agreement with Alnylam Pharmaceuticals, consults for Flagship Pioneering and Chroma Medicines, and serves on the SAB of Ubiquitix. These interests are not directly related to the contents of this paper.

## Extended Data Figure Legends

**Extended Data Fig. 1.**
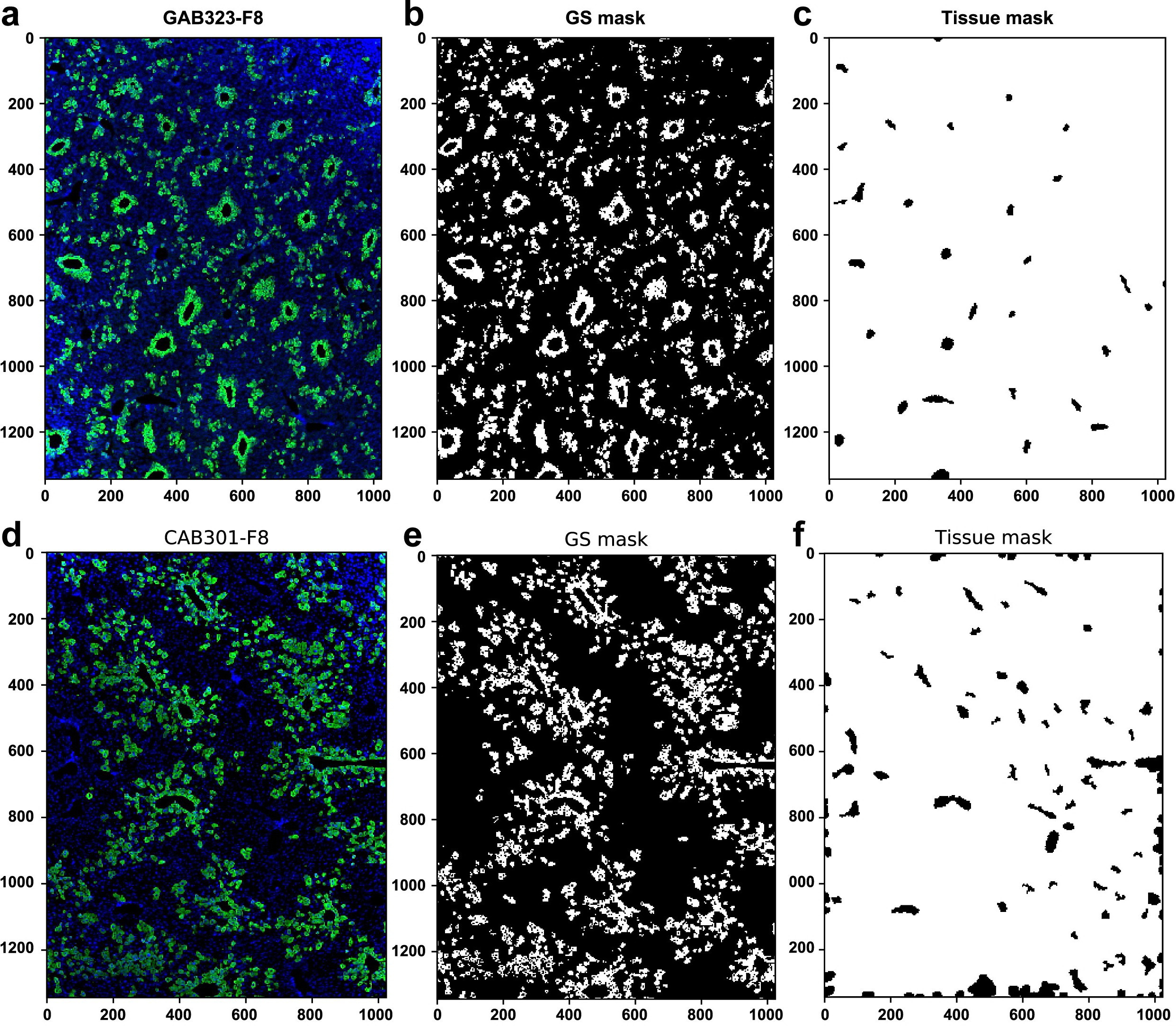
An image analysis algorithm to measure GS+ tissue area. **a.** Original composite IF image of a Zone 1 *Gls2-CreERT2; Ctnnb1^(ex3)^ ^fl/+^; Arid2^fl/fl^* liver stained for GS and DAPI at 2 months post-tamoxifen. **b.** A mask is segmented from the GS channel (GS mask). **c.** The DAPI channel is used to generate the tissue mask. Dividing the number of pixels in the GS mask by the number of pixels in the DAPI mask gives a proportion of the image that is GS+. **d-f.** Original composite IF image, GS mask, and tissue mask of a zone 3 *Cyp1a2-CreERT2; Ctnnb1^(ex3)^ ^fl/+^; Arid2^fl/fl^* mouse at 3 months post-tamoxifen.

**Extended Data Fig. 2.**
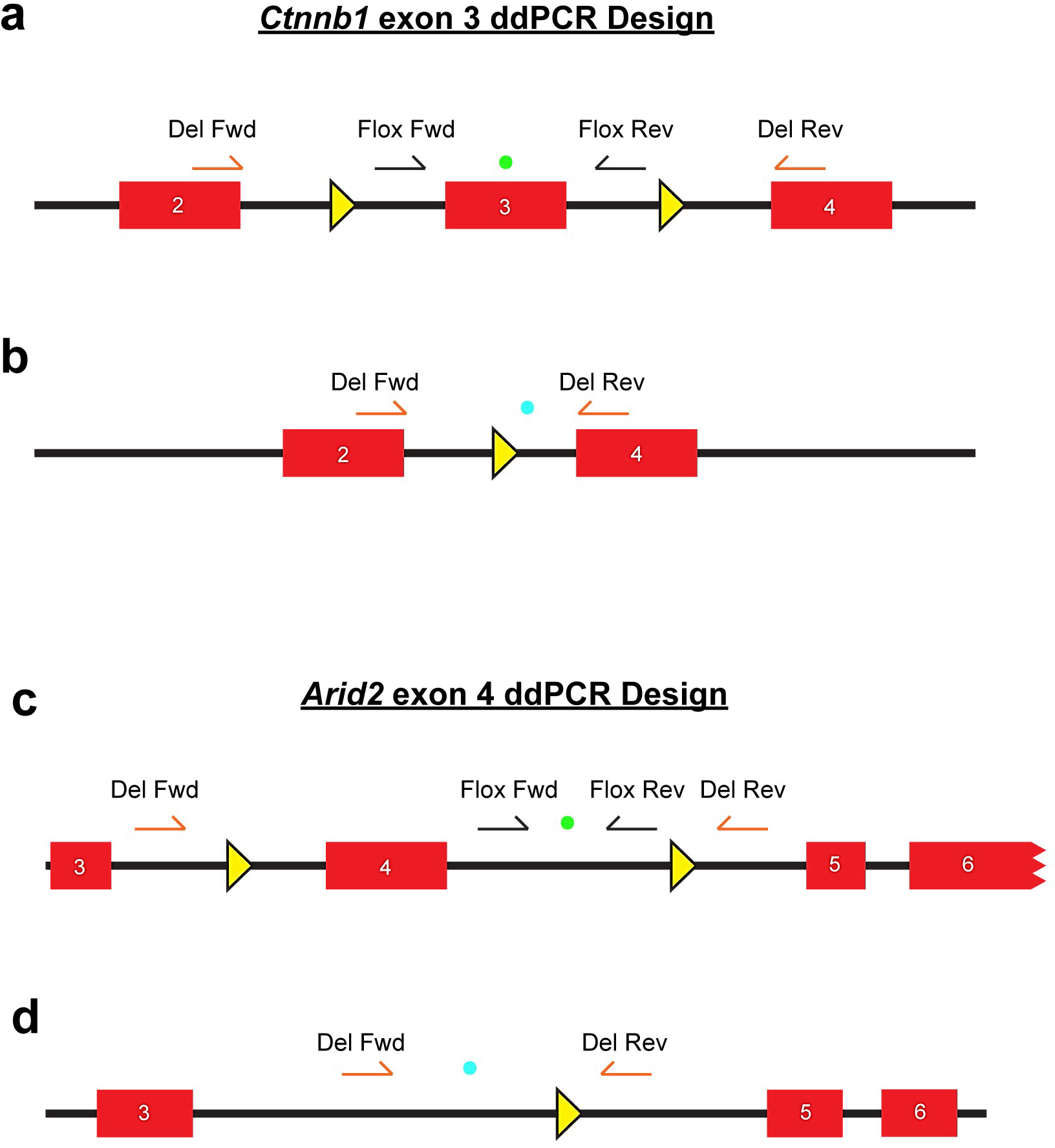
A ddPCR assay to measure *Ctnnb1* and *Arid2* mutant alleles. **a.** A primer probe set was designed for the *Ctnnb1^(ex3)/flox^* allele. The primers and probe (HEX) are entirely contained within the floxed region. **b.** Another primer probe set was designed for the *Ctnnb1^(ex3)/Δ^* allele. The primers and probe (FAM) are outside of the floxed region. **c.** A primer probe set was designed for the *Arid2^fl/fl^* allele. The primers and probe (HEX) are entirely contained within the floxed region. **d.** Another primer probe set was designed for the *Arid2^Δ/Δ^* allele. The primers and probe (FAM) are outside of the floxed region.

**Extended Data Fig. 3.**
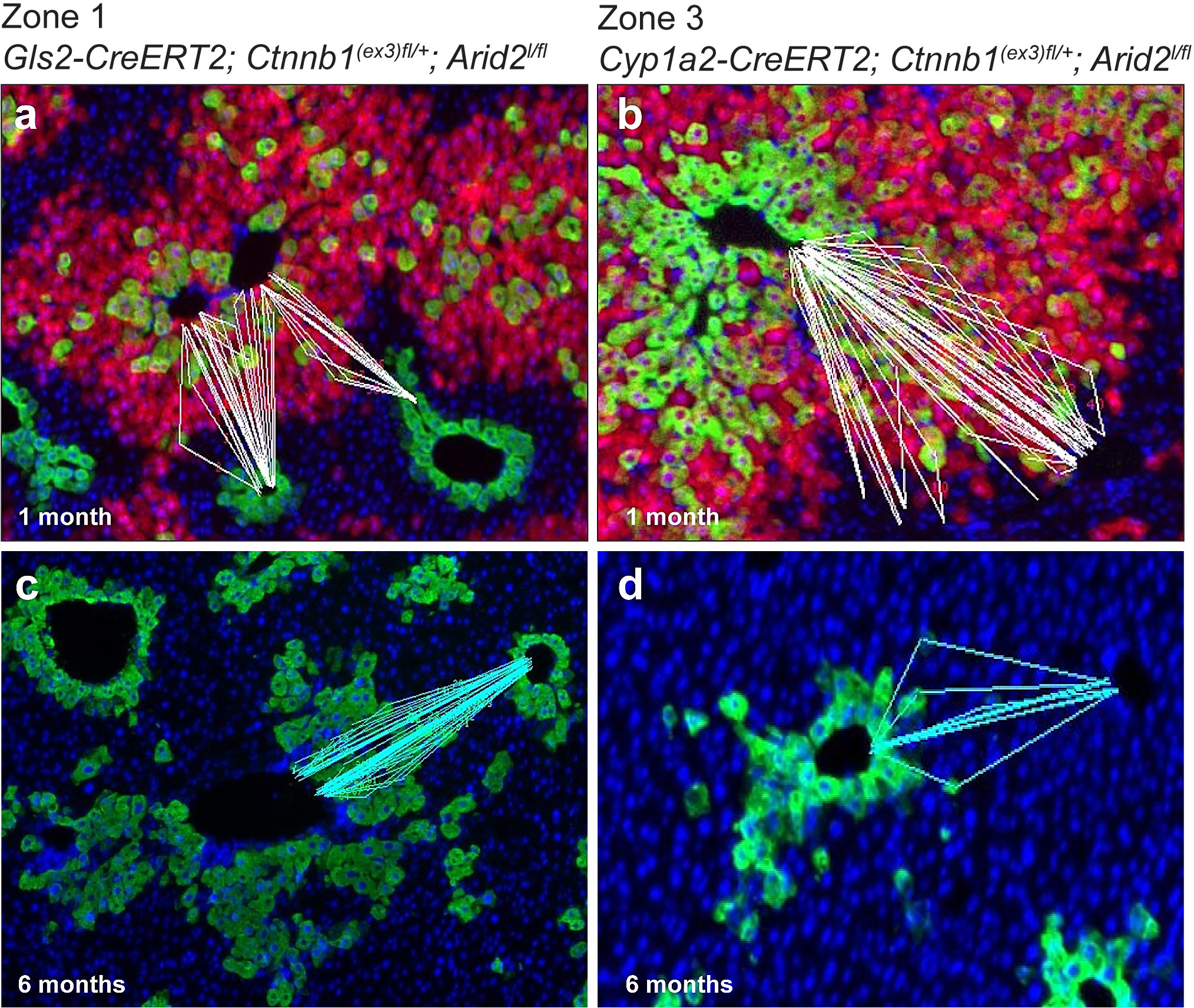
Determining the CV-PT position of mutant hepatocytes. **a,b.** The position index (P.I.) was calculated by measuring and recording the distances from a cell to the nearest CV and PT, and the distance between the CV and PT. Examples from a 1 month **(a)** zone 1 *Gls2-CreERT2; Ctnnb1^(ex3)^ ^fl/+^; Arid2^fl/fl^* liver and **(b)** zone 3 *Cyp1a2-CreERT2; Ctnnb1^(ex3)^ ^fl/+^; Arid2^fl/fl^* liver are shown. **c,d.** Examples from 6 month **(c)** zone 1 liver and **(d)** zone 3 liver are shown.

**Extended Data Fig. 4.**
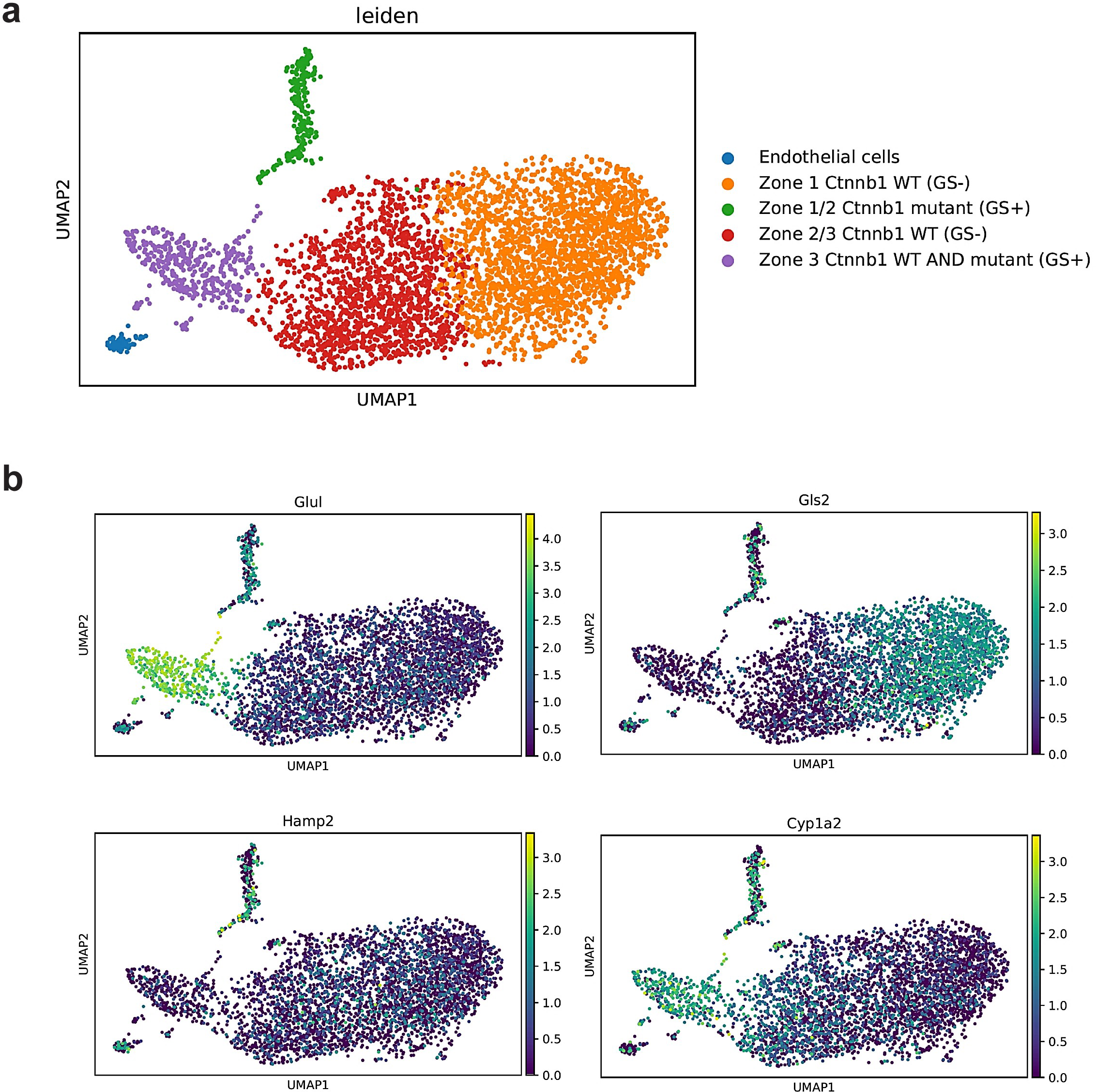
snRNA-seq reveals that the identity and state of mutant hepatocytes depends on their location within the lobule. **a,b.** From the five cell populations identified through leiden clustering of snRNA-seq of a zone 3 liver, Zone 1 *Ctnnb1* WT hepatocytes are identified by enriched expression of zone 1 genes such as *Gls2* and no expression of *Glul* (GS). Zone 3 *Ctnnb1* mutant and WT hepatocytes are identified by enriched expression of zone 3 genes such as *Glul* and *Cyp1a2.* A population of zone 1/2 *Ctnnb1* mutant hepatocytes is identified by enriched expression of *Gls2, Hamp2*, and *Glul*.

**Extended Data Fig. 5.**
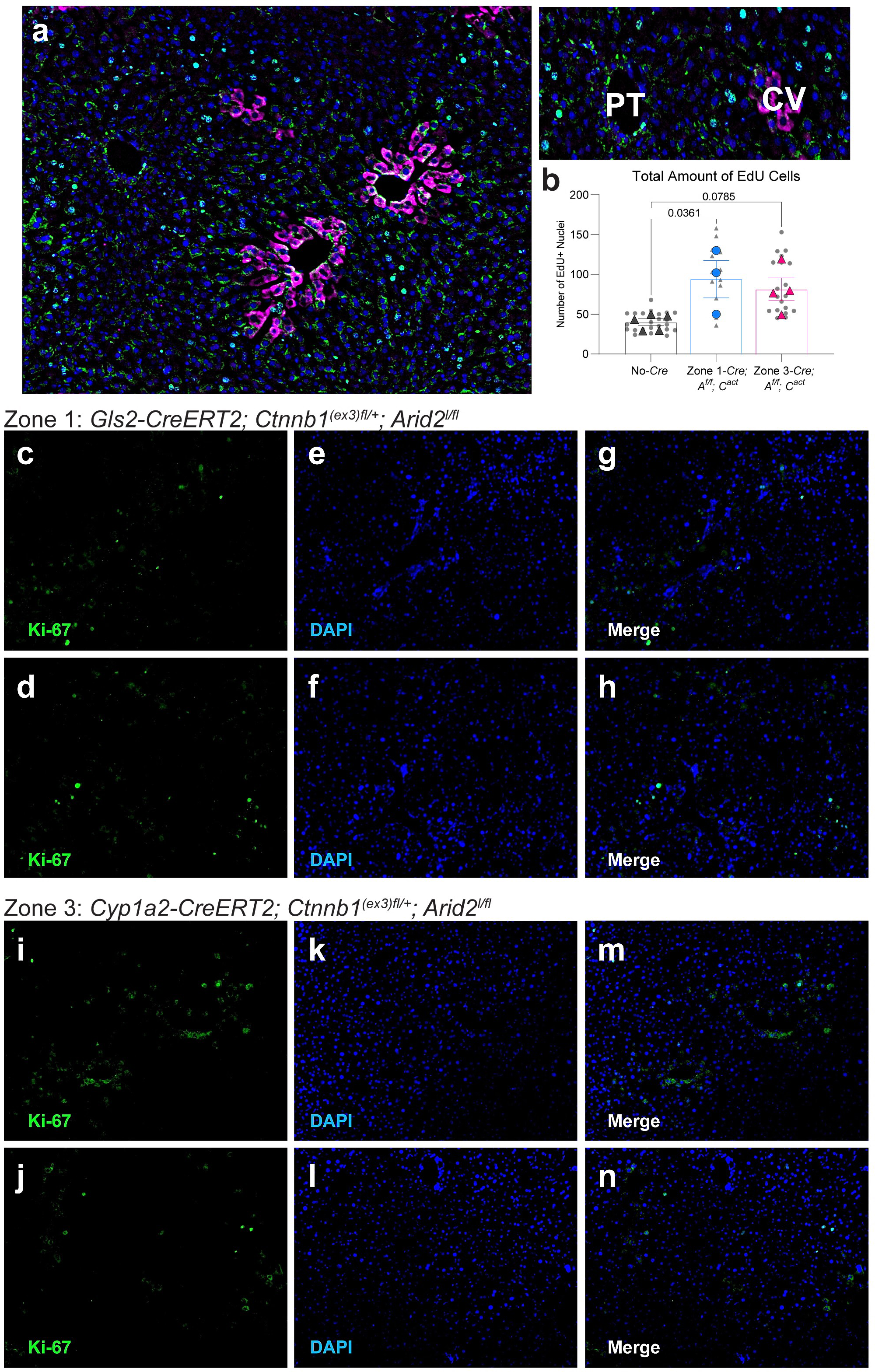
There is more cell proliferation in zone 1 mutant livers than in zone 3 mutant livers. **a.** Control mice without Cre were given EdU water 1 month after tamoxifen and their livers were collected after 10 days. A PT-to-CV axis is shown with EdU+ nuclei distribution. **b.** Total absolute number of EdU+ cells were measured from liver images. Smaller black dots represent counts from single 10X images, and larger colored dots represent counts of each mouse averaged from at least 4 or more 10X images. EdU pulse chase experiments were performed on two different times and the data was aggregated. **c-h.** Two representative zone 1 *Gls2-CreERT2; Ctnnb1^(ex3)^ ^fl/+^; Arid2^fl/fl^* livers stained for Ki-67 6 months after tamoxifen. **(c)** and **(d)** are examples of Ki-67 staining. **(e)** and **(f)** are examples of nuclear DAPI staining. **(g)** and **(h)** are merged images. **i-n.** Two representative zone 3 *Cyp1a2-CreERT2; Ctnnb1^(ex3)^ ^fl/+^; Arid2^fl/fl^* livers stained for Ki-67 6 months after tamoxifen. **(i)** and **(j)** are examples of Ki-67 staining. **(k)** and **(l)** are examples of nuclear DAPI staining. **(m)** and **(n)** are merged images. Data in bar graphs are displayed as mean ± SEM, and statistical analyses were performed using a one-way ANOVA ( **b**).

**Extended Data Fig. 6.**
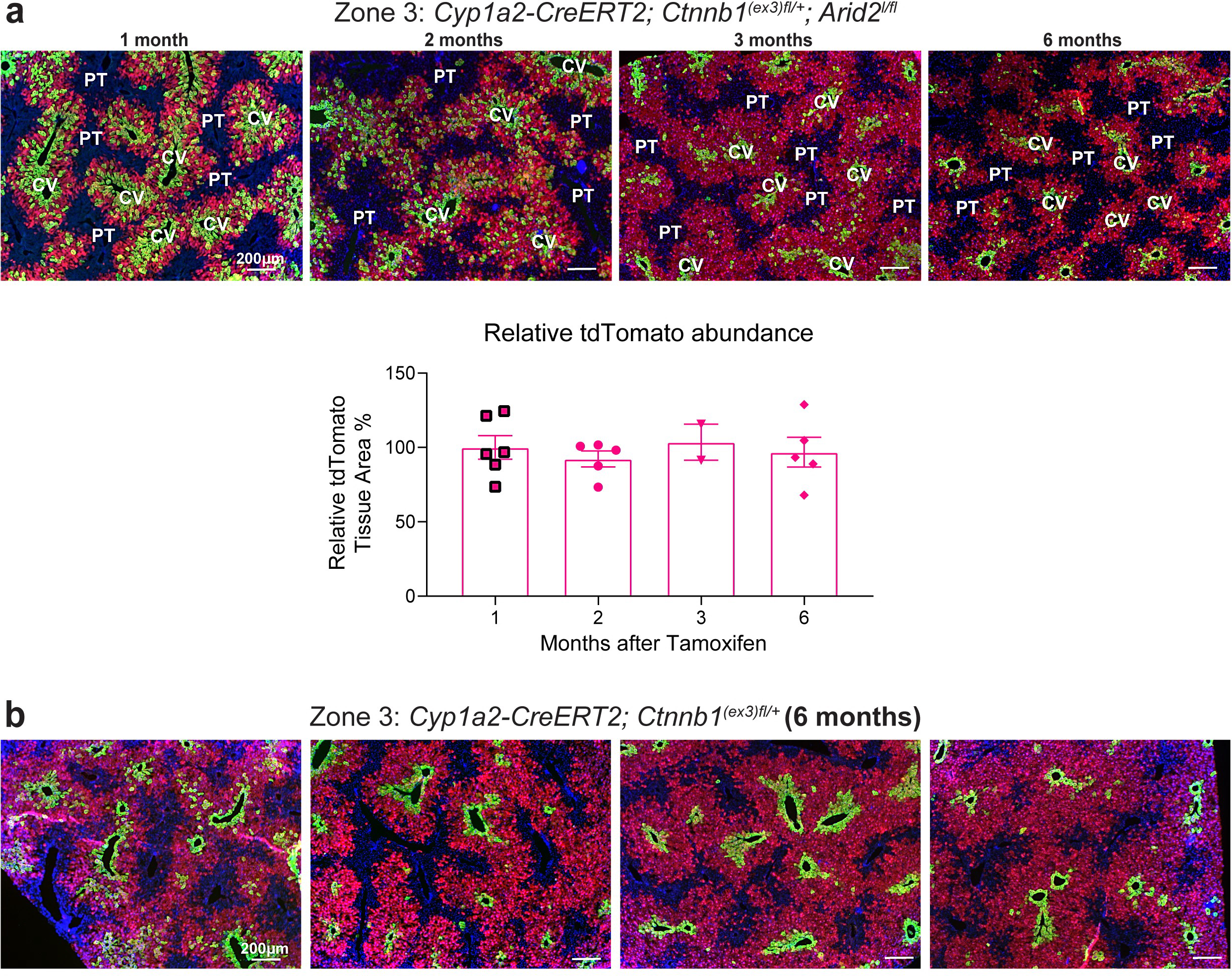
Lineage tracing with tdTomato area shows decreased proportion of GS+ mutants. **a.** GS-stained livers from zone 3 *Cyp1a2-CreERT2; Ctnnb1^(ex3)^ ^fl/+^; Arid2^fl/fl^; Rosa26^LSL-tdTomato/+^* mice from 1 to 6 months after tamoxifen. tdTomato area remains constant while the GS+ mutant clone population decreases in number. The bar graph shows tdTomato+ area percentages for zone 3 mice over time with data normalized to the 1 month time point, which was designated as 100%. For tdTomato quantification in zone 3 mice at 1, 2, 3, and 6 months, each dot represents one mouse and n = 6, 5, 2, 5 mice. **b.** GS-stained livers from zone 3 *Cyp1a2-CreERT2; Ctnnb1^(ex3)^ ^fl/+^* single mutant mice at 6 months after tamoxifen. Each panel represents one liver from one mouse. The aged single mutant mice show a similar reduction in GS+ mutant clones as the aged double mutant mice at the same time points. Data in bar graphs are displayed as mean ± SEM.

**Extended Data Fig. 7.**
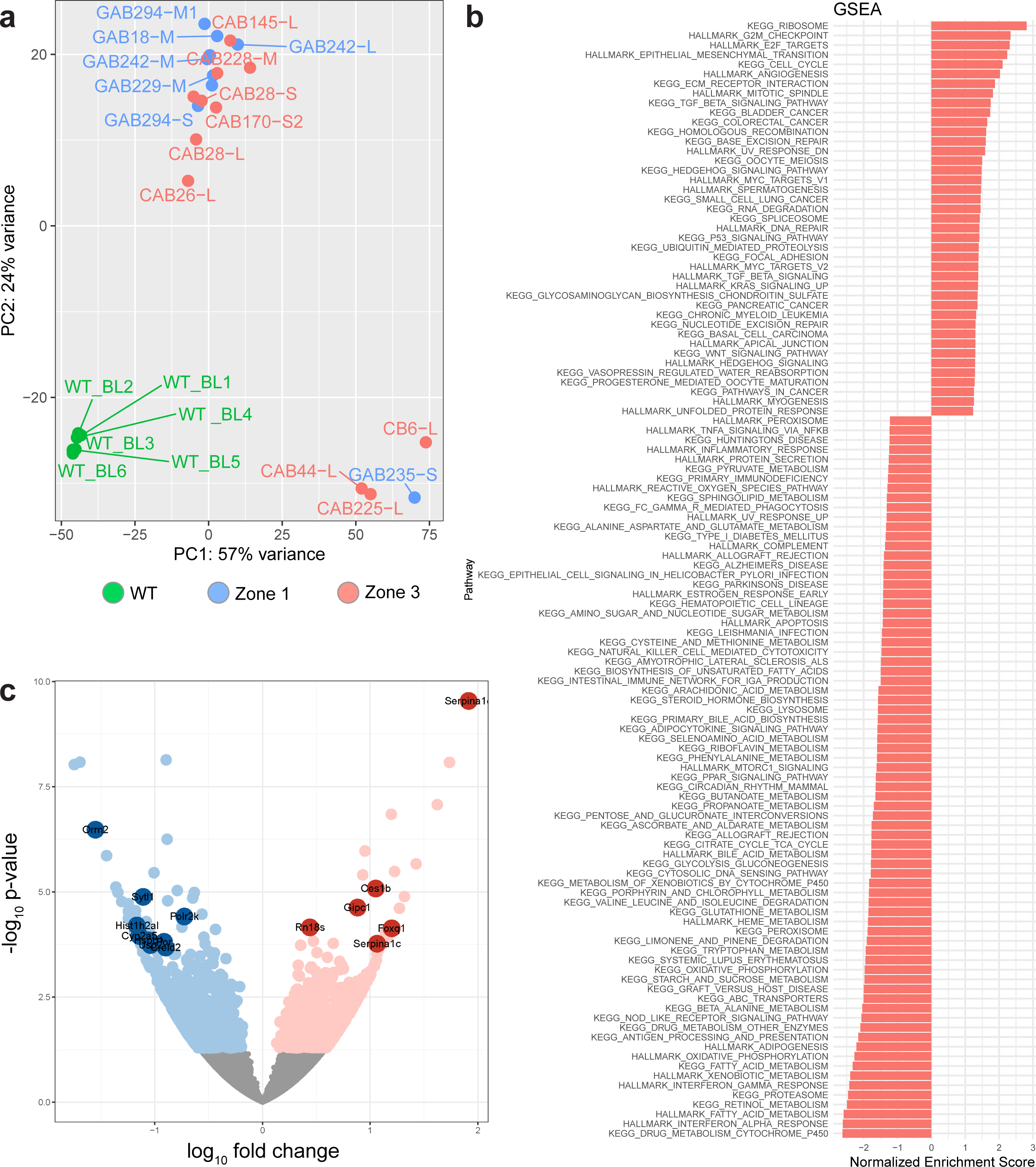
Characterization of *Ctnnb1* mutant, *Arid2* deleted tumors originating from zone 1 vs. zone 3. **a.** Principal components analysis score plot for WT livers (n=6), zone 1 tumors (n=9), and zone 3 tumors (n=11) analyzed in bulk RNA-seq. Both types of zone tumors are different from the WT livers, but for the most part, tumors tend to group together. **b.** Gene set enrichment analysis for pathways in zone 3 tumors relative to zone 1 tumors shows enrichment of cell cycle and cancer growth-related genes. **c.** Volcano plot displaying the log_10_(fold change) of zone 3 tumors relative to zone 1 tumors on the x axis and the -log_10_(p value) on the y axis. Bulk RNA-seq in tumors from zone 1 (n = 9) and zone 3 (n = 10) revealed 67 differentially expressed genes, including some that are normally zonated in the liver (highlighted genes). Details of the statistical analysis for the RNA-seq can be found in the Methods section.

**Extended Data Fig. 8.**
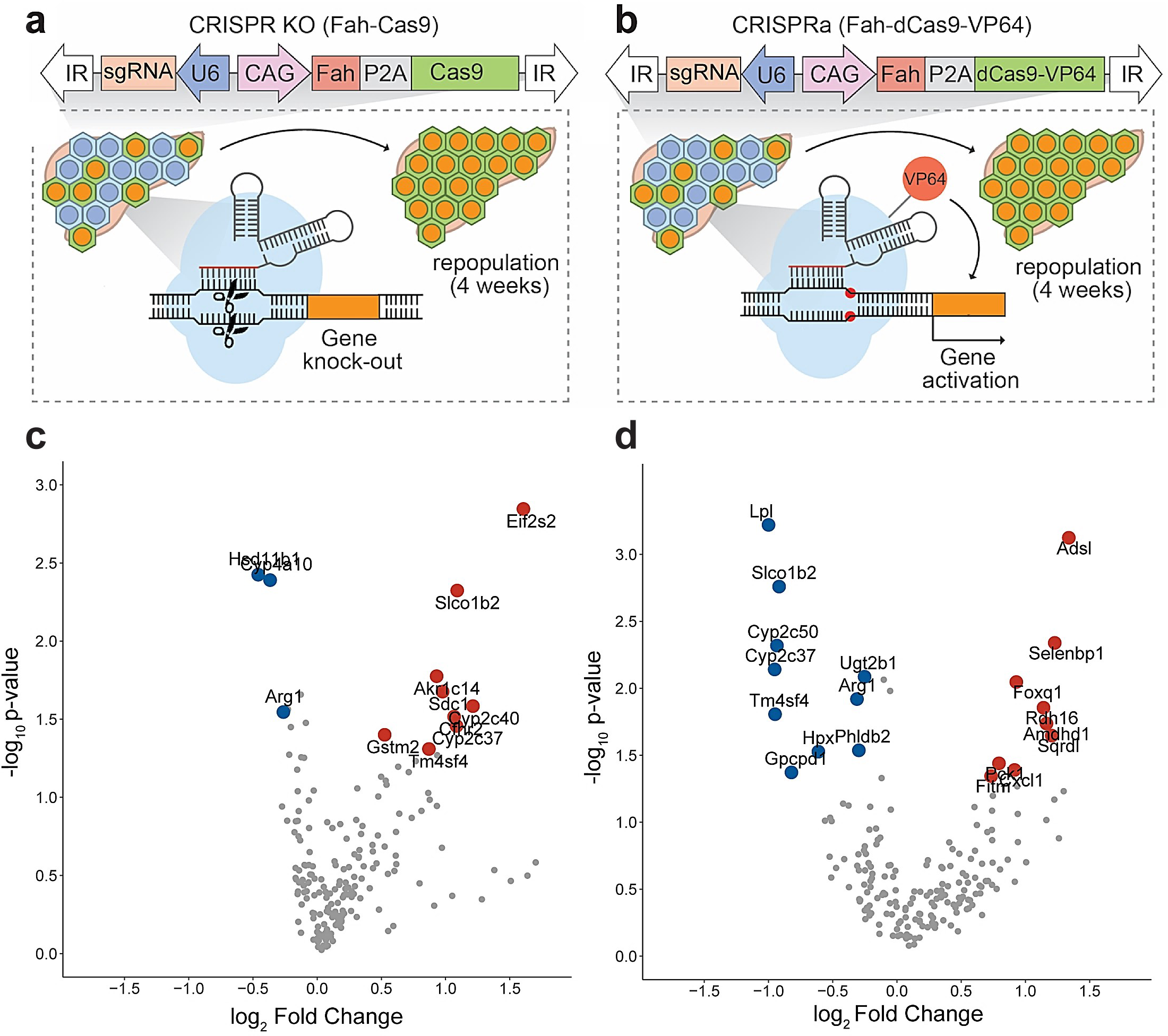
An *in vivo* CRISPR screen identifies zonated genes that mediate differences in hepatocyte proliferation. **a,b.** *In vivo* **(a)** CRISPR KO and **(b)** CRISPRa screening strategies were used to identify zonated genes that could mediate mutant hepatocyte fate. Briefly, sgRNA libraries were cloned into a transposon carrying the gene for Cas9 or dCas9-VP64 and Fah. Transposons + SB100 transposase were hydrodynamically injected into *Fah^-/-^* mice (n = 10 for KO screen, n = 7 for activation screen). The mice were taken off of NTBC-supplemented water, allowing for transfected hepatocytes to repopulate the liver over a period of 4 weeks. Then, genomic DNA was collected from the livers of the mice, and the integrated sgRNAs were amplified and sequenced. **c,d.** Several candidate genes were identified in **(c)** the KO screen and **(d)** the CRISPRa screen. sgRNA representation was analyzed by the MAGeCK algorithm using the default settings.

**Extended Data Fig. 9.**
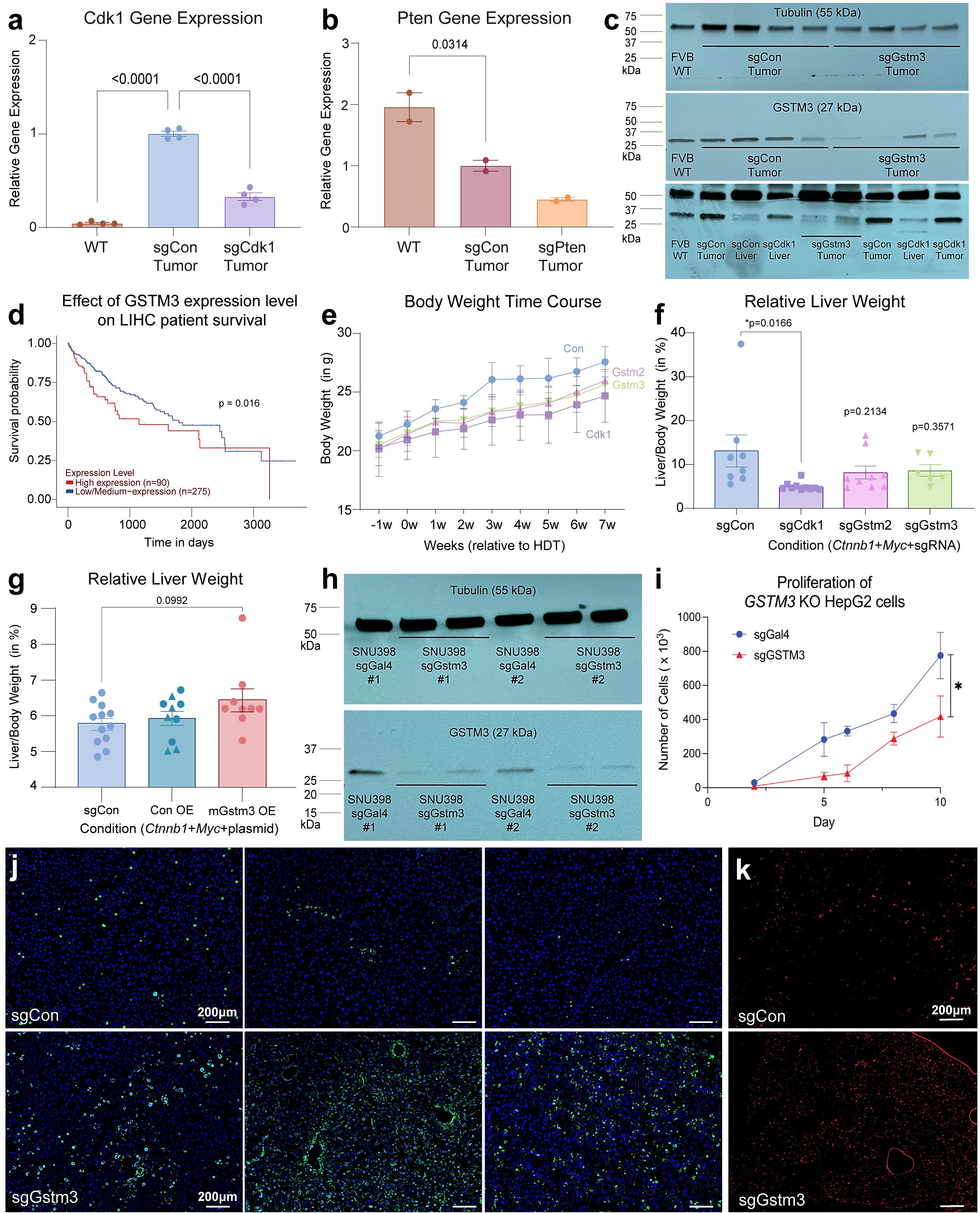
Additional data for the HDT tumor models, related to Fig. 4. **a,b.** RT-qPCR for HDT CRISPR KO controls *Cdk1* (**a**) and *Pten* (**b**). mRNA expression levels are expressed relative to sgCon tumor tissue, which is designated as 1. Each dot represents one liver simple from one mouse for Cdk1 (n=4 mice) and Pten (n=2). **c.** Western blot image of β-tubulin and GSTM3 expression in WT FVB livers and HDT sgCon, sgCdk1, and sgGstm3 livers or tumors. Top and center panel show one blot probed for GSTM3 (center), stripped, and reprobed for β-tubulin (top). Bottom panel shows full blot for partial lanes shown in Fig. 4 and is probed for both GSTM3 (27 kDa) and β-tubulin (55 kDa). The sgCon and sgGstm3 lanes from Fig. 4 are vertically reflected from the original blot to mirror the order of the RT-qPCR results. Each lane contains lysate from one liver from one mouse. **d.** UALCAN analysis of *GSTM3* expression and how it correlates with survival of HCC patients. **e.** BW tracked over time until 7 weeks after HDT in survival C57BL/6J mice from Fig. 4e. For sgCon, sgCdk1, sgGstm2, and sgGstm3 groups, n = 9, 8, 10, 9 mice. **f.** LW/BW at 5 weeks after HDT in C57BL/6J mice. Each dot represents one mouse, and for sgCon, sgCdk1, sgGstm2, and sgGstm3 groups, n = 8, 10, 9, 6 mice. **g.** LW/BW at 1 week after HDT, with GFP (triangle) and Luciferase (square) overexpression groups as controls along with sgCon compared to GSTM3 overexpression mice. Each symbol represents one mouse, and for sgCon, Con OE, and Gstm3 OE groups, n = 12, 10, 9 mice. Overexpression HDT experiments were performed two independent times, and data was aggregated. **h.** Western blot image of β-tubulin and GSTM3 expression in transduced *Gstm3* KO SNU398 cells or control *Gal4* KO cells to validate lentiviral sgRNA efficiency. Top and bottom panel show one blot probed for GSTM3 (bottom), stripped, and reprobed for β-tubulin (top). Each lane contains lysate from one liver from one mouse. Cells were transduced two independent times, and samples from each experiment are indicated by the number. **i.** Proliferation time course of HepG2 cells with lentiviral KO of *Gstm3* or control KO of *Gal4* measuring cell number in days after seeding. Experiments were performed three independent times, and data was aggregated. **j.** Additional TUNEL-stained livers from HDT sgCon (top row) and sgGstm3 (bottom row) mice 1 week after injection. Each image is from one mouse. **k.** Superoxide ROS levels visualized through DHE staining for sgCon and sgGstm3 livers 1 week after injection. Each image is from one mouse. Data in bar graphs are displayed as mean ± SEM, and statistical analyses were performed using a one-way ANOVA when comparing multiple HDT groups ( **a,b,f,g**) or a two-tailed and unpaired Student’s t-test when comparing two groups (**i**).

## Notes

### Competing Interest Statement

The authors have declared no competing interest.

